# Conformational changes in the Niemann-Pick Type C1 protein NCR1 drive sterol translocation

**DOI:** 10.1101/2023.09.08.556848

**Authors:** Kelly M. Frain, Emil Dedic, Lynette Nel, Anastasiia Bohush, Esben Olesen, David L. Stokes, Bjørn Panyella Pedersen

**Affiliations:** Department of Molecular Biology and Genetics, Aarhus University, Universitetsbyen 81, Aarhus C 8000, Denmark; Aarhus Institute of Advanced Studies, Aarhus University, Høegh-Guldbergs Gade 6B, DK-8000 Aarhus C, Denmark; Skirball Institute of Biomolecular Medicine, Department of Cell Biology, New York University, Grossman School of Medicine, New York, NY 10016, United States

**Keywords:** sterol uptake, ergosterol, cholesterol, vacuole, lysosome, Niemann-Pick type C protein, NPC system, NCR1, NPC1, cryo-EM, glycocalyx

## Abstract

The membrane protein Niemann-Pick Type C1 protein (NPC1, named NCR1 in yeast) is central to sterol homeostasis in eukaryotes. *Saccharomyces cerevisiae* NCR1 is localized to the vacuolar membrane, where it is suggested to carry sterols across the protective glycocalyx and deposit them into the vacuolar membrane. However, documentation of a vacuolar glycocalyx in fungi is lacking and the mechanism for sterol translocation has remained unclear. Here we provide evidence that a glycocalyx is indeed present inside isolated *Saccharomyces cerevisiae* vacuoles, and report four cryo-EM structures of NCR1 in two distinct conformations that elucidate how it moves sterol through the glycocalyx. The two conformations, named “tense” and “relaxed”, illustrate movement of sterol through a tunnel formed by the luminal domains. Based on these structures and on comparison with other members of the Resistance-Nodulation-Division (RND) superfamily we propose a transport model that links changes in the luminal domains with a cycle of protonation and deprotonation within the transmembrane region of the protein. Our model suggests that NPC proteins work by a generalized RND mechanism where the transmembrane domains form a ’motor-unit’ that sequentially adopts a tense and relaxed conformation to drive changes in luminal/extracellular domains.

**SIGNIFICANCE STATEMENT:** Niemann-Pick Type C1 (NPC1, named NCR1 in yeast) proteins play a critical role in sterol homeostasis by facilitating the integration of sterols into membranes of acidic organelles like lysosomes and vacuoles. The inner surface of these organelles’ membranes is shielded by the glycocalyx. Here, we provide evidence that a glycocalyx is present in vacuoles from *Saccharomyces cerevisiae* and demonstrate that NCR1 transports sterols across it by undergoing conformational changes. Our structures suggest a transport model where sterol transport is linked to proton-driven changes in the transmembrane region. This work sheds light on the mechanism of NPC1 protein function and has broad implications for understanding lysosomal storage disorders and for mechanisms employed by members of the Resistance-Nodulation-Division (RND) superfamily.

## MAIN TEXT

## INTRODUCTION

Niemann Pick Type C1 (NPC1) membrane proteins are critical for establishing cell sterol homeostasis in eukaryotes. Sterols are typically transported via the endocytic pathway to acidic organelles (lysosomes and vacuoles) before integration via NPC1 into the membrane for further redistribution (Winkler et al., 2022). Failure to integrate sterols into the membrane leads to lipid accumulation in lysosomes and in humans gives rise to a neurodegenerative lysosomal storage disorder called NPC disease (Evans & Hendriksz, 2017; Ribeiro et al., 2001). The *Saccharomyces cerevisiae* ortholog, named NCR1, has been used as a model system to understand NPC disease and mechanisms associated with sterol uptake in general (Berger et al., 2005; Malathi et al., 2004; Sturley, 2000; Winkler et al., 2019; Winkler et al., 2022).

A central role of NCR1 and NPC1 is proposed to be the transport of sterols across the glycocalyx, a polysaccharide coating that maintains the integrity of the vacuolar/lysosomal membrane (Fig. 1A). While the presence of a glycocalyx is well documented in the mammalian lysosome, the presumption that it also exists in vacuoles, the fungal equivalent of the lysosome, is not supported by direct evidence (Lehle et al., 2006; Neiss, 1984; Tarbell & Cancel, 2016). Nevertheless, three steps have been delineated for the membrane integration of sterols by NPC proteins: loading, transfer and transport (Winkler et al., 2022). Loading refers to the delivery of sterol to a luminal N-terminal domain (NTD) of NCR1. Thereafter, sterol is transferred from the NTD binding pocket into a ∼60-Å long tunnel that spans the glycocalyx. This tunnel is formed at the interface between a middle luminal domain (MLD) and a C-terminal domain (CTD) (Fig. 1B) (Damke et al., 2010; Winkler et al., 2019). Transport refers to the movement of sterols through this tunnel before their release into the luminal leaflet of the membrane (Winkler et al., 2019; Winkler et al., 2022). The membrane domain has 13 transmembrane helices (M1-M13) divided into the sterol-sensing domain (SSD, M2-M7), and the pseudo-SSD (pSSD, M8-M13), and sterol release after transport happens through a gate in the SSD at the end of the tunnel. Two-fold pseudo-symmetry relates the SSD to the pSSD and the MLD to the CTD (Fig. 1B).

**Figure 1.**
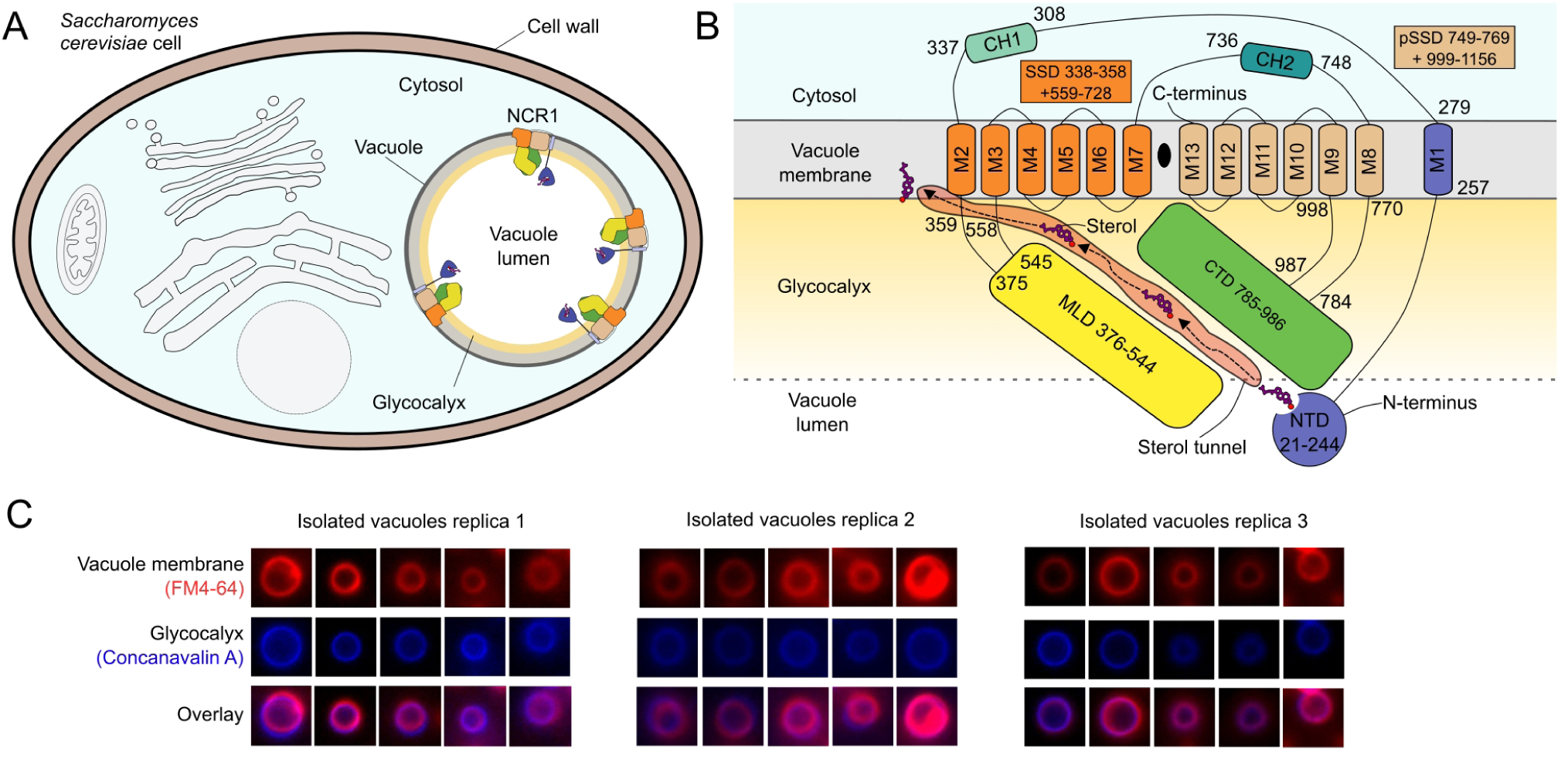
Cellular localization of NCR1, its topology and the presence of the vacuolar glycocalyx. **(A)** *S. cerevisiae* cell organization, focusing on the vacuole organelle inside. Within the acidic vacuole resides the membrane protein NCR1. **(B)** Topology schematic of NCR1. The path of sterol movement is shown from the NTD, translocating through the tunnel and integrated into the vacuolar membrane. **(C)** Enlarged pictures of isolated vacuoles from three replicates. Vacuolar membranes were stained red with lipophilic dye FM4-64 and the glycocalyx was stained blue with Concanavalin A conjugated to an AlexaFluor 350.

NCR1 belongs to the Resistance-Nodulation-Division (RND) superfamily (Nikaido, 2018). The RND superfamily contains nine families that are widespread in prokaryotes and eukaryotes and that transport a variety of compounds (Klenotic et al., 2021). Most RND family members are not “transporters” in the regular sense since these proteins do not move substrate from one side of the membrane to the other, but rather insert or extract substrate from the membrane or periplasmic space (Klenotic et al., 2021). This process is best characterized by the multi-drug efflux pump, acriflavine resistance protein B (AcrB) (Nikaido, 2018; Zwama & Yamaguchi, 2018). AcrB couples substrate transport through a porter domain (topologically equivalent to the MLD and CTD of NCR1) to the proton motive force as mediated by a network of titratable residues at the rotational pseudo-symmetry point of the SSD and pSSD, and a variety of RND family members have been implicated more broadly in coupling transport to electrochemical membrane potential at this location (Adams et al., 2021; Furukawa et al., 2018; Moseng et al., 2021; Tam et al., 2021; Wang et al., 2021; Zhang et al., 2018; Zwama & Yamaguchi, 2018). A similar network is found in NPC proteins centered on a conserved acidic pair: Asp631 and Glu1068 in NCR1 (Winkler et al., 2019). This conserved acidic pair have been shown to be essential for function of NCR1 and NPC1 (Qian et al., 2020; Winkler et al., 2019; Zhang et al., 2017). In analogy to AcrB, it has been proposed that protons might provide the driving force for sterol movement through NPC by harnessing the pH gradient across lysosomal and vacuolar membranes (Winkler et al., 2022). Structures of human NPC1 at pH 5.5 and pH 8.0 have indeed revealed distinct, pH-dependent conformations, but these structural changes were ascribed to a pH-dependent auto-inhibitory mechanism which was proposed to suppress activity prior to arriving in the lysosome/vacuole (Qian et al., 2020). A mechanistic model for sterol transport through the tunnel, as well as energy requirements, including possible coupling to the transmembrane proton gradient, remain unclear. It is also unclear if NTD loading and transfer in the NPC proteins are rigidly coupled to their transport cycle, or whether the NTD functions independently of the RND core domains (SSD + MLD and pSSD + CTD) (Winkler et al., 2022).

Here we provide evidence that the glycocalyx is present in yeast vacuoles, confirming a defining role of NCR1 as a glycocalyx bypass for sterols. To further explore the roles of protons in sterol transport, we solved cryo-EM structures of NCR1 at pH 5.5 and pH 7.5. The structures show that the NTD of NCR1 is not involved in the sterol transport mechanism, but serves as a tethered binding domain to receive substrate and transfer it to the tunnel. The structures reveal distinct conformational changes that are consistent with two key states of a cycle driven by protonation and deprotonation of the acidic pair. A series of residues link the acidic pair to the SSD gate and coordinates inter-domain movements between the MLD and the CTD, driving sterol movement through the tunnel towards the membrane.

## RESULTS

### A glycocalyx is present in the yeast vacuole

To establish the presence of a glycocalyx on the vacuolar membrane, we isolated intact vacuoles from *S. cerevisiae* by first removing the cell wall with zymolyase and then using differential centrifugation through a Ficoll gradient. This preparation was stained with FM4-64, a lipophilic dye routinely used for visualizing vacuole morphology, and the glycocalyx was stained with Concanavalin A, a lectin that selectively binds α-mannopyranosyl and α-glucopyranosy glycoproteins, conjugated to an AlexaFluor 350. Fig. 1C shows vesicles in the expected size range for vacuoles of 2-5 μm, many of which displayed co-localization of lipid and saccharide staining, indicating the presence of a polysaccharide matrix. These observations were consistent across three replicates involving examination of 150 individual vesicles, though only ∼25% showed Concanavalin A staining (Fig. S1), indicating either a preponderance of non-vacuolar vesicles in the preparation (e.g., from other components of the endomembrane system) and/or difficulty in stain penetration as is required since the glycocalyx would be residing on the inner surface of the vacuole. These results are consistent with the presence of a glycocalyx on the vacuolar membrane which, to our knowledge, has not been previously shown.

### Structures of NCR1 show two key conformations

To assess the role of protons in sterol transport, we examined whether NCR1 adopted different conformations at high and low concentrations of protons. Specifically, we utilized pH 5.5 (high proton concentration) and pH 7.5 (low proton concentration) to mimic high and low protonation states of NCR1, respectively, and solved structures of NCR1 under these conditions (Fig. 2A, 2B and Fig. S2-S5 and Table S1).

**Figure 2.**
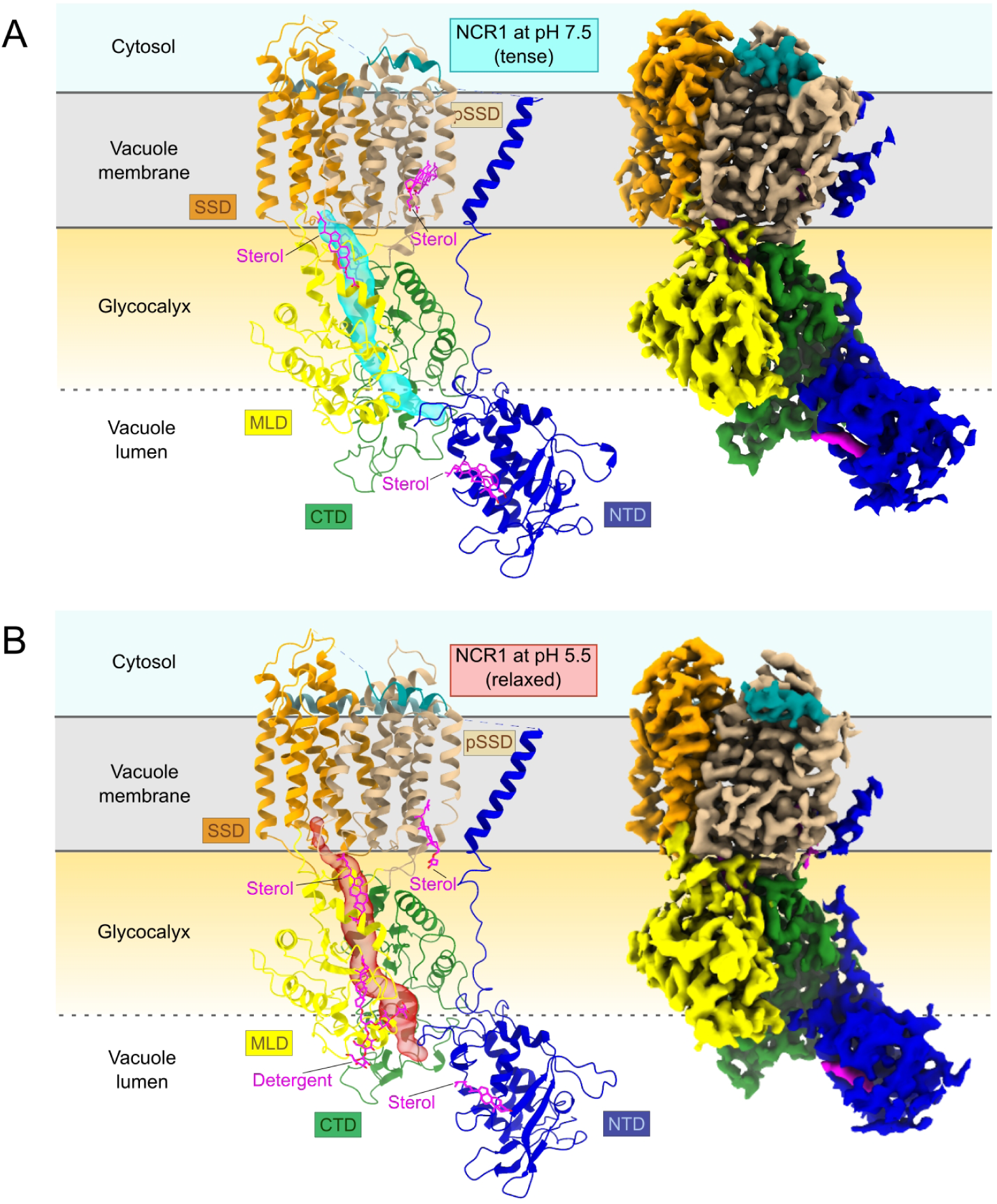
Comparison of NCR1 structures at pH 7.5 and pH 5.5. **(A)** Model and map of NCR1 in GDN detergent at pH 7.5. A tunnel (cyan) stretches from the NTD to the SSD. Ligands are depicted in magenta and include an ergosterol in the NTD, a CHS in the tunnel and a CHS in a hydrophobic patch between the pSSD and M1. **(B)** Model and map of NCR1 in GDN detergent at pH 5.5. A tunnel (red) stretches from the NTD to the SSD. Ligands are depicted in magenta and include an ergosterol in the NTD, a GDN detergent molecule in the tunnel, a CHS also in the tunnel and a CHS in a hydrophobic patch between the pSSD and M1.

We solved the structure of NCR1 at pH 7.5 using Glyco-diosgenin (GDN) as the detergent at a resolution of 3.3 Å (Fig. 2A, Fig 3A, 3B and Fig. S2). In this structure, an ergosterol is present in the NTD and two cholesterol hemmisuccinate (CHS) molecules are found: one in the tunnel between the MLD and CTD and one at a hydrophobic patch between M1 and pSSD (Fig 3A, 3B, Fig. S6). The presence of CHS likely reflects the addition of this amphiphile during initial solubilization of the cell membranes whereas ergosterol was likely bound *in vivo* and was co-purified together with NCR1. Notably, the conformation revealed by this structure is similar to that observed in the previously published crystal structure of NCR1 (PDB code 6R4L) (Winkler et al., 2019).

**Figure 3.**
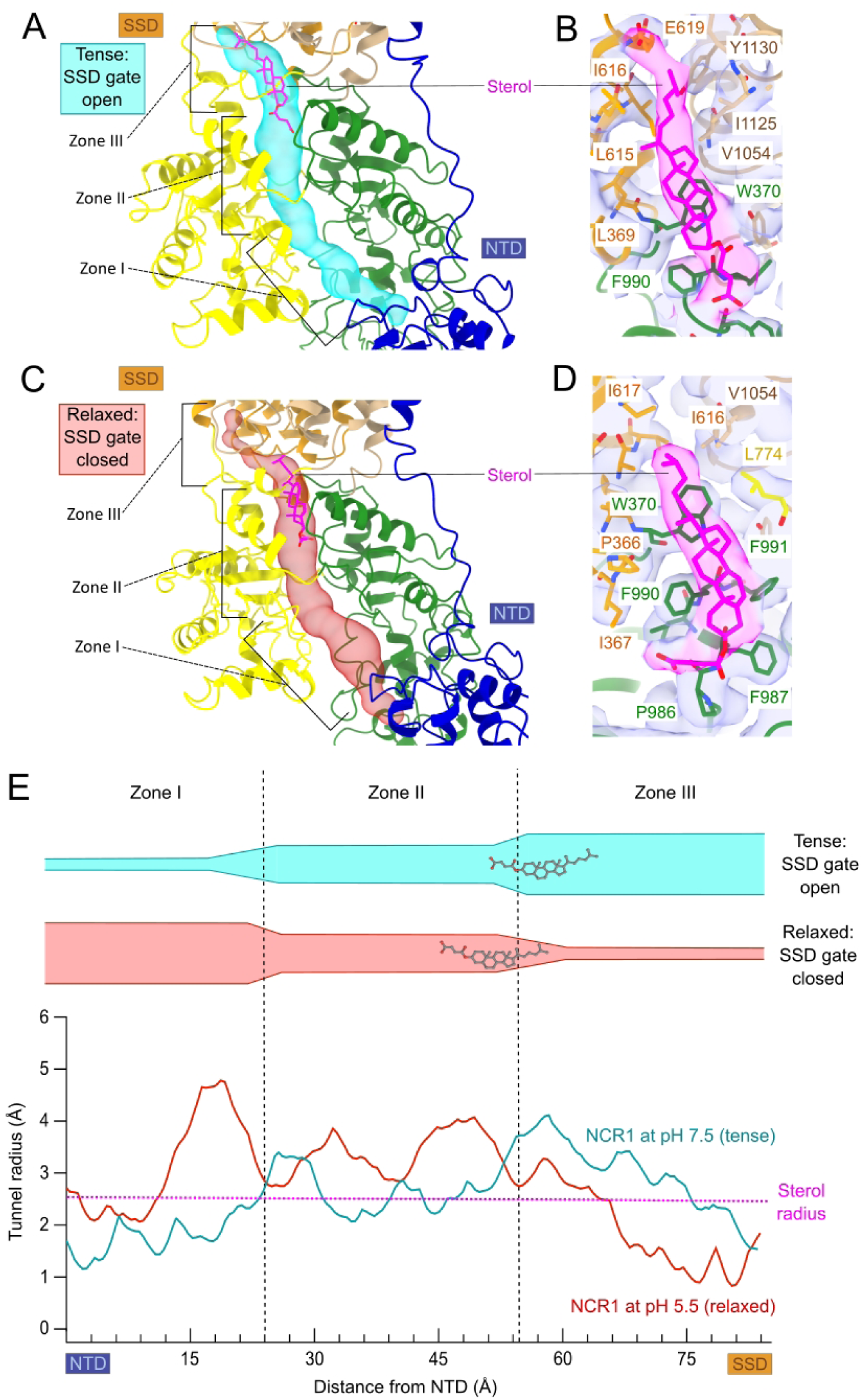
NCR1 Tunnel profile comparison between pH 7.5 and pH 5.5. **(A)** Zoomed model of NCR1 at pH 7.5 tunnel (cyan) between the MLD (yellow) and the CTD (green). Zones I, II and III of the tunnel are indicated with brackets. **(B)** Density of CHS ligand in NCR1 in GDN at pH 7.5. **(C)** Zoomed model of NCR1 pH 5.5 tunnel (red) between the MLD (yellow) and the CTD (green). Zones I, II and III of the tunnel are indicated with brackets. **(D)** Density of CHS ligand in NCR1 in GDN at pH 5.5. **(E)** Graph indicating tunnel radius (Å) versus distance from the NTD (Å) for the tunnel in NCR1 at pH 7.5 (cyan) and pH 5.5 (red). Sterol radius (2.5 Å) is shown with a dotted magenta line along the X axis. The graph is divided into Zones, I, II and III, which section the schematic representation of the tunnels, including the sterol location, shown above the graph.

We then proceeded to solve the structure at pH 5.5 in GDN, also at a resolution of 3.3 Å (Fig. 2B, Fig 3C, 3D and Fig. S3). In this second structure, a GDN detergent molecule was seen in addition to the two CHS ligands (Fig 3C, 3D, Fig. S6). The GDN molecule is located at the bottom of the tunnel near the NTD. One CHS is located in the tunnel near the membrane surface - displaced compared to the first structure - while the other CHS molecule occupies the same hydrophobic patch between M1 and the pSSD.

Comparison of the two structures at pH 7.5 and 5.5 reveals a series of key differences, making it clear that the structure at pH 5.5 displays a different conformation compared to the structure at pH 7.5. When considered individually, RMSD values comparing MLD, CTD, SSD and pSSD are minimal: 0.6 Å for CTD and 0.9 Å for MLD, 1.2 Å for SSD and 1.0 for pSSD. However, rigid body movements between the domains are apparent in the context of the entire molecule. In the transmembrane region, superposition of the SSD reveals a ∼2.5 Å rigid-body displacement of the pSSD. That is, in the pH 5.5 structure, the SSD and pSSD move away from each other and adopt a distinct conformation we name the ’relaxed’ conformation. In contrast, the SSD and pSSD are closer together at pH 7.5, which we name the ’tense’ conformation (Figure S7A and Table 1). There is also a movement of the CTD relative to the MLD: after structural alignment of the CTD, there is a ∼3 Å displacement and slight rotation of the MLD in the two conformations (Fig. S7B).

**Table 1.** Summary of the conformations of NCR1 solved by X-ray crystallography and cryo-EM.

| Conformation | Tense |  | Relaxed |  |  |
| --- | --- | --- | --- | --- | --- |
| SSD gate | Open |  | Closed |  |  |
| Acidic pair network | D631-E1068-H1072 |  | D631-E1068-K1111 |  |  |
| Sterol location | zone III | zone I | zone II | zone II | zone II |
| Condition | pH 7.5<br>GDN | pH 6.1<br>LCP | pH 5.5<br>GDN | pH 5.5<br>LMNG | pH 7.5<br>Peptidisc |
| PDB | 8QEB | 6R4L | 8QEC | 8QED | 8QEE |

Because GDN occupied the tunnel at pH 5.5, we solved an additional structure in Lauryl Maltose Neopentyl Glycol (LMNG) detergent, also at pH 5.5 and also at 3.3 Å resolution, to rule out an influence of this detergent on the observed conformation (Fig. S4 and Fig. S8A). This third structure is devoid of GDN or other detergent molecules that could influence the conformation, but still retains the sterols found in the other two structures (Fig. S6). It is in the relaxed conformation and essentially identical to the structure in GDN at pH 5.5 (RMSD of 0.8 Å) and the acidic pair is in the same position (Fig. S8B), indicating that the GDN detergent molecule is not the determinant of the new NCR1 conformation.

In all the structures, the ligand positions are well-defined. However, both here and in previous work on NCR1 and human NPC1, the pose of ligands in the tunnel have been ambiguous due to limited resolution. An amphipathic helix solubilization system, called peptidisc, has been shown to lock proteins in defined conformations, often improving resolution (Carlson et al., 2018; Ung et al., 2022). Indeed, use of peptidisc improved the resolution to 2.4 Å (Fig. S5, Fig. S9A), thus enabling us to confirm the pose of sterol within the tunnel, with the hydroxyl group oriented toward the lumen of the vacuole (Fig. S9B) as modeled in the X-ray structure and as suggested earlier (Kwon et al., 2009). In addition, water molecules are visible in a cavity within the membrane domain near the acid pair (Fig. S9C). However, this structure adopts the relaxed conformation seen at pH 5.5, even though it was produced at pH 7.5 and the NTD and M1 helix were not visible at any stage during image processing. In parallel we solved an additional structure at pH 7.5 in LMNG to 3.3 Å that also displayed a similar behavior as the peptidisc sample: no density of the NTD or M1, and an observed relaxed conformation of the transmembrane SSD and pSSD domains (date not shown). This suggests that the lipid and detergent environment influence the ability of the M1 helix and the NTD to associate to the RND core and can also restrict movements of the RND core domains. Innate flexibility of the NTD and M1 helix is consistent with the lower resolution of these elements compared to the RND core in all our structures, consistent with loose association with the core. In fact, in all of our datasets we identified a sub-population of particles that produced structures lacking the NTD domain and M1 helix (Fig. S2E, S3E, S4E). Flexibility of the NTD has been suggested to assist in loading of the sterol, and it is noteworthy that, in the current maps, ordering of the NTD had no effect on the RND core domains, which displayed no structural shifts when structures from the subpopulations (with and without NTD) were compared.

### Changes in the tunnel shape drive sterol transport

The dimensions of the tunnel, formed by the MLD and CTD, undergo substantial changes as a result of the conformational shifts between these two domains. From the two structures determined in GDN, we characterize the tunnel dimensions extending from the NTD (near Asp557) to the beginning of the SSD gate (near Tyr1130) (Fig. 3A, 3C). From the resulting tunnel radius profiles, we define three distinct zones: Zone I near the NTD, Zone II in the middle of the MLD and CTD and Zone III near the SSD gate. In the relaxed conformation, Zone I is broad with the radius peaking at 4.8 Å, while the same zone becomes narrow with an average radius around 2 Å in the tense conformation (Fig. 3E). Zone III displays inverse behavior with the tunnel narrowing considerably to an average radius of 2 Å in the relaxed conformation, whereas the tunnel widens to ∼3 Å radius in the tense conformation. In Zone II, the differences between the two conformations are minimal with an average radius of ∼ 3 Å. A broader comparison of NCR1 structures indicates that tunnel profiles strongly correlate with the conformation of the transmembrane domain: the tunnel profile for NCR1 at pH 7.5 is consistent with that from the X-ray structure and both are in the tense conformation, whereas structures in the relaxed conformation have the opposite tunnel radius profile (Fig. S10). This correlation indicates allosteric coupling between the membrane domain (tense vs. relaxed) and the MLD and CLD, which determine the tunnel shape. Furthermore, these comparisons suggest that concerted expansion and contraction of the tunnel is linked to the tense and relaxed conformation of the transmembrane domain and potentially represents a mechanism for propelling the sterol through the tunnel. This mechanism is consistent with changes in the position of the sterol observed within the tunnel. Specifically, in the relaxed conformation, there is well defined density for sterol at the boundary of Zones II and III (Fig. 3C, 3D), whereas sterol density from the tense conformation is displaced ∼6 Å towards the SSD gate and the density elongated (Fig. 3A, 3B), indicating mobility of the sterol as it gets closer to the tunnel exit.

### The acidic pair network links tense and relaxed conformations to sterol transport

A network of interactions at the pseudosymmetry axis relating SSD to pSSD govern transition between the tense and relaxed conformations. This network centers on the conserved acidic pair, Asp631 (M5) and Glu1068 (M11), which alternately interact with His1072 (M11) and Lys1111 (M12) (Fig. 4A). In particular, the acidic pair coordinate His1072 (distance < 3.5 Å) in the tense conformation (NCR1 at pH 7.5), but in the relaxed conformation (NCR1 at pH 5.5), the acidic pair pivots downward to form a new interaction with Lys1111 (also <3.5 Å Fig. 4B). During this transition, Lys1111 undergoes the most dramatic movement, being buried between M3 and M5 in the tense conformation, but swinging towards the acidic pair in the relaxed conformation thus drawing them away from His1072 (Fig. 4C). The bonding network between Asp631, Glu1068 and His1072 in the tense conformation induces tighter packing between SSD and pSSD, whereas this interface relaxes and the domains move apart as a result of the new Asp631-Glu1068-Lys1111 network that characterizes the relaxed conformation. This relaxation creates a small cavity between the SSD and pSSD, which is occupied by a number of water molecules visible in the higher resolution peptidisc structure (Fig. S9C).

**Figure 4.**
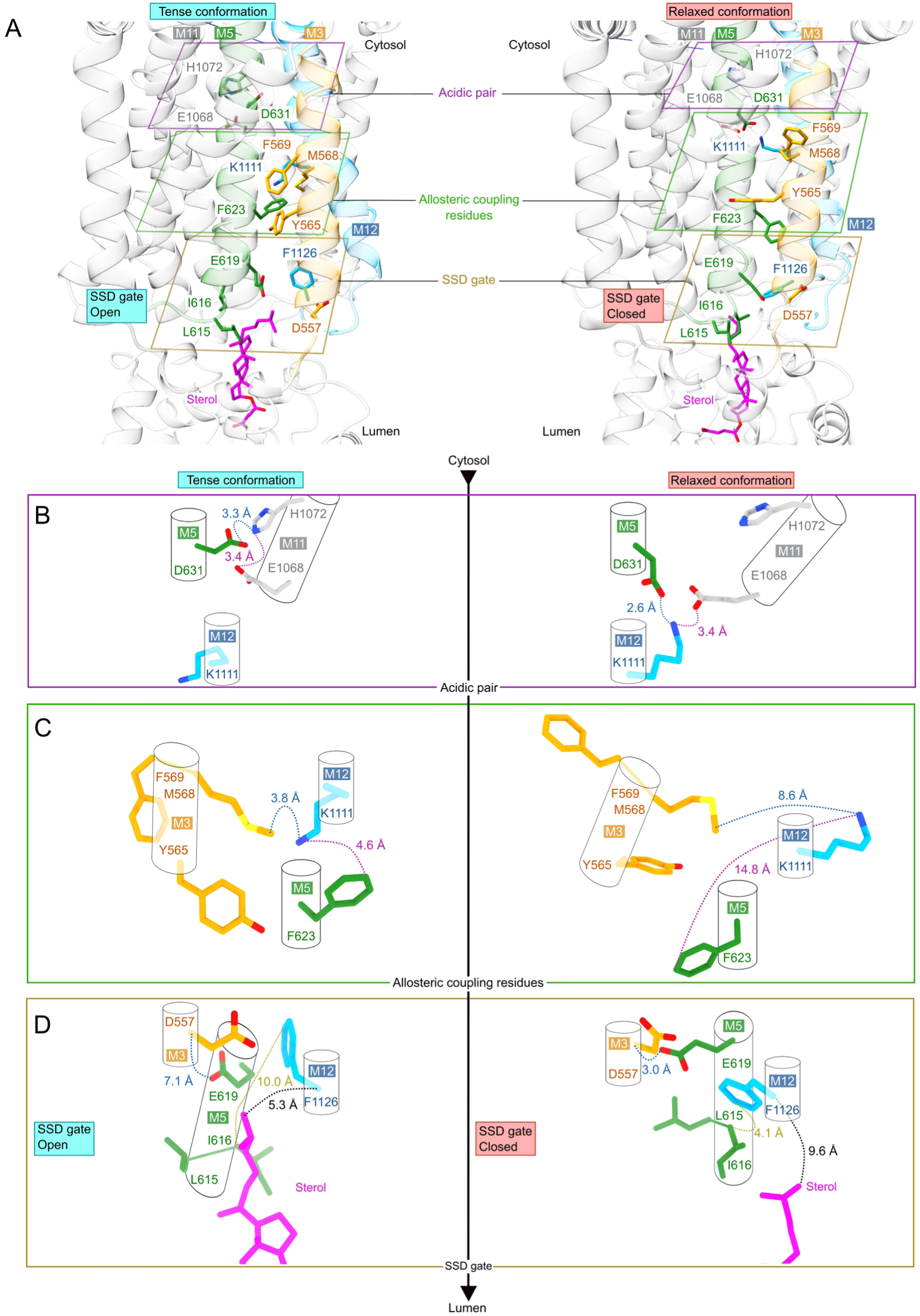
Acidic pair and allosteric coupling residues controlling the opening and closing of the SSD gate. **(A)** Side view of the transmembrane domains in the tense (left) and relaxed (right) conformation. The models are transparent with the relevant atoms shown as non-transparent sticks. Helices of interest are colored orange (M3), green (M5) and blue (M12). Residues of interest are grouped by sectioning planes: the acidic pair (purple box), allosteric coupling residues (green box) and the SSD gate (yellow box). **(B)** The acidic pair and interacting residues of the tense (left) and relaxed (right) conformation. **(C)** Allosteric coupling residues of helices M3 and M5. **(D)** Key residues of the SSD gate.

The transition between these two states is allosterically coupled to the SSD gate and the conformation of the tunnel, thus controlling integration of sterol into the membrane. M5 undergoes displacement during this transition making it a plausible element for allosterically linking the acidic pair to the opening or closing of the SSD gate (Fig. 4C). Specifically, the SSD gate is governed by movements at the luminal ends of M3, M5, M12 and the associated loops. In going from relaxed to tense conformations, the distance between Glu619 (M5) and Asp557 (M3) increases from 3.0 Å to 7.1 Å and the distance between Ile616 and Phe1126 increases from 4.1 Å to 10 Å (Fig. 4D). These changes are due to a dramatic swing of the Phe1126 sidechain as well as reconfiguration of the M4/M5 loop and the first turn of M5 and serve to control passage of sterol - with a diameter of ∼5 Å through the SSD gate. Operation of the SSD gate is also correlated with widening and narrowing of Zone III of the tunnel, thus promoting sterol transport: as the gate opens in the relaxed conformation, the sterol molecule in the tunnel moves ∼6 Å toward the membrane with its aliphatic tail protruding through the gate and ready for integration into the membrane (Fig. 4D).

## DISCUSSION

In lysosomes, low pH combines with hydrolytic enzymes to break down metabolites for recycling. In yeast and plants, the vacuole fulfills this function in addition to playing an expanded role in storage of nutrients and ions (Tan et al., 2019; Van Ho et al., 2002; Winkler et al., 2019; Winkler et al., 2022). For the lysosome, it is well-established that the inner leaflet of the membrane is protected by the glycocalyx against autodigestion (Kosicek et al., 2018; Liu et al., 2020). It is therefore reasonable to expect a similar barrier for the inner membrane of vacuoles. Here, we demonstrate co-localization of lipid-and saccharide-stain, thus supporting the presence of a glycocalyx in yeast vacuoles as a shared feature of degradative organelles.

An unresolved question for the function of NPC proteins is whether loading and transport of sterol are coupled or whether loading of the NTD is independent of subsequent transport through the tunnel (Winkler et al., 2022). Our data suggests the latter: that the NTD acts as a tethered binding domain with no discernible coupling to the transport process. Specifically, comparison of structures with and without the NTD shows that its association with the other luminal domains has no effect on their conformation. Moreover, the presence of M1 has no effect on the membrane SSD and pSSD domains. In all our structures, the NTD is loaded with ergosterol and oriented in a post-loading state, despite pH-dependent conformational changes to the RND core. Indeed, analyses of *npc* genes in Eukaryotes revealed that some fungal genomes encode the NTD as a separate gene product that is not covalently tethered to the RND core, reinforcing the importance of the NTD for sterol delivery, but not necessitating it to be part of one protein chain to accomplish transport (Adebali et al., 2016). Furthermore, M1 and the NTD are not features found in related RND protein-families which appear to employ analogous transport mechanisms (Zhang et al., 2017). Earlier studies of NPC1ΔNTD also support this idea by showing the isolated NTD can transfer substrate to the core domains (Trinh et al., 2018). Nevertheless, tethering of the NTD would serve to increase local sterol concentration and thereby increase the efficiency and specificity of the transfer process (Dyla et al., 2022). Here it is also noteworthy that the association of the NTD and M1 and the movements of the SSD and pSSD appears highly dependent on the lipid/detergent environment. For future work it would be relevant to use replace detergents with e.g. a lipid/nanodisc system to better mimic the physiological lipid bilayer environment.

The high resolution of the NCR1 structure solved in peptidisc offers valuable insights into the orientation of sterol and clearly shows that the polar head group of the sterol faces the vacuolar lumen (Fig. S9B). This pose supports the hand-off model for the NTD loading step, which positions the sterol to be transferred into the tunnel in the optimal orientation for integration into the inner leaflet of the organellar membrane (Kwon et al., 2009).

Our analysis of two different conformations of NCR1 indicate that the tunnel changes its profile due to rigid body movement of the MLD relative to the CTD (Fig. 3A, 3C and Fig. S7B). These changes are accompanied by a shift in the sterol position within the tunnel. Similar conformational changes have been observed in other RND proteins and they are proposed to generate a peristaltic mode of transport in the multi-drug efflux transporter AcrB (Seeger et al., 2006; Zwama & Yamaguchi, 2018). In this respect, it is notable that NPC1 function was abolished when cysteine crosslinking was used to prevent interdomain movements of MLD and CTD (Saha et al., 2020). Thus, it seems that these changes in the tunnel are coordinated with opening and closing of the SSD gate in NCR1.

For NPC1, pH-dependent conformational change has been ascribed to an inhibitory mechanism with the high pH form representing an inactive state, adopted while the protein is trafficked through the endomembrane system (Qian et al., 2020). According to this hypothesis, the low pH form represents a constitutively active state achieved within the lysosome. However, the disposition of the SSD gate in both human NPC1 and now in our data on yeast NCR1 are inconsistent with this scenario. Specifically, the SSD gate is open at high pH, when the protein is suggested to be inactive, and closed at low pH, when the protein is suggested to be active. Alternatively, the energy coupling mechanism employed by AcrB and other RND superfamily members and our observations of sterol displacement in the tunnel between two distinct conformations suggest a different interpretation of the observed conformational change.

We propose a model to explain the observed conformations of NPC proteins, where the tense and relaxed conformations represent two key conformations in a proton-driven sterol transport cycle (Fig. 5). In this model, the acidic pair Asp631 and Glu1068 in NCR1 forms a network analogous to AcrB (Fig. S11) at the interface of the SSD and pSSD. Depending on the protonation state of the network - in our case induced by pH of the sample - interactions are made with either His1072 or Lys1111. Thus, our model invokes a cycle with Lys1111 being the initial proton acceptor from the lumen in the tense conformation (Fig. 5, step 1). When Lys1111 accepts a proton, perhaps via Glu619 and Tyr565, it swings toward the acidic pair, inducing the relaxed conformation. As a result, the SSD gate closes and there is a coordinated narrowing of Zone III and widening of Zone I of the tunnel (Fig. 5, step 2). Formation of the tripartite network between the acidic pair and Lys1111 induces His1072 to release a proton to the cytosol (Fig. 5, step 3). Asp631 and Glu1068 now engage in a 3-step transfer of the proton from Lys1111 to His1072 (Fig. 5, step 3-5). After donating its proton, Lys1111 swings back into the initial position, nestled into a hydrophobic pocket between M3 and M5, while the acidic pair changes conformation to interact with His1072 to generate the tense conformation. Together, these changes lead to a new conformational shift of the MLD and CTD, driving peristaltic movements of sterol within the tunnel and opening the SSD gate (Fig. 5, step 1). Binding of a new luminal proton to Lys1111 initiates a new cycle of peristaltic movement.

**Figure 5.**
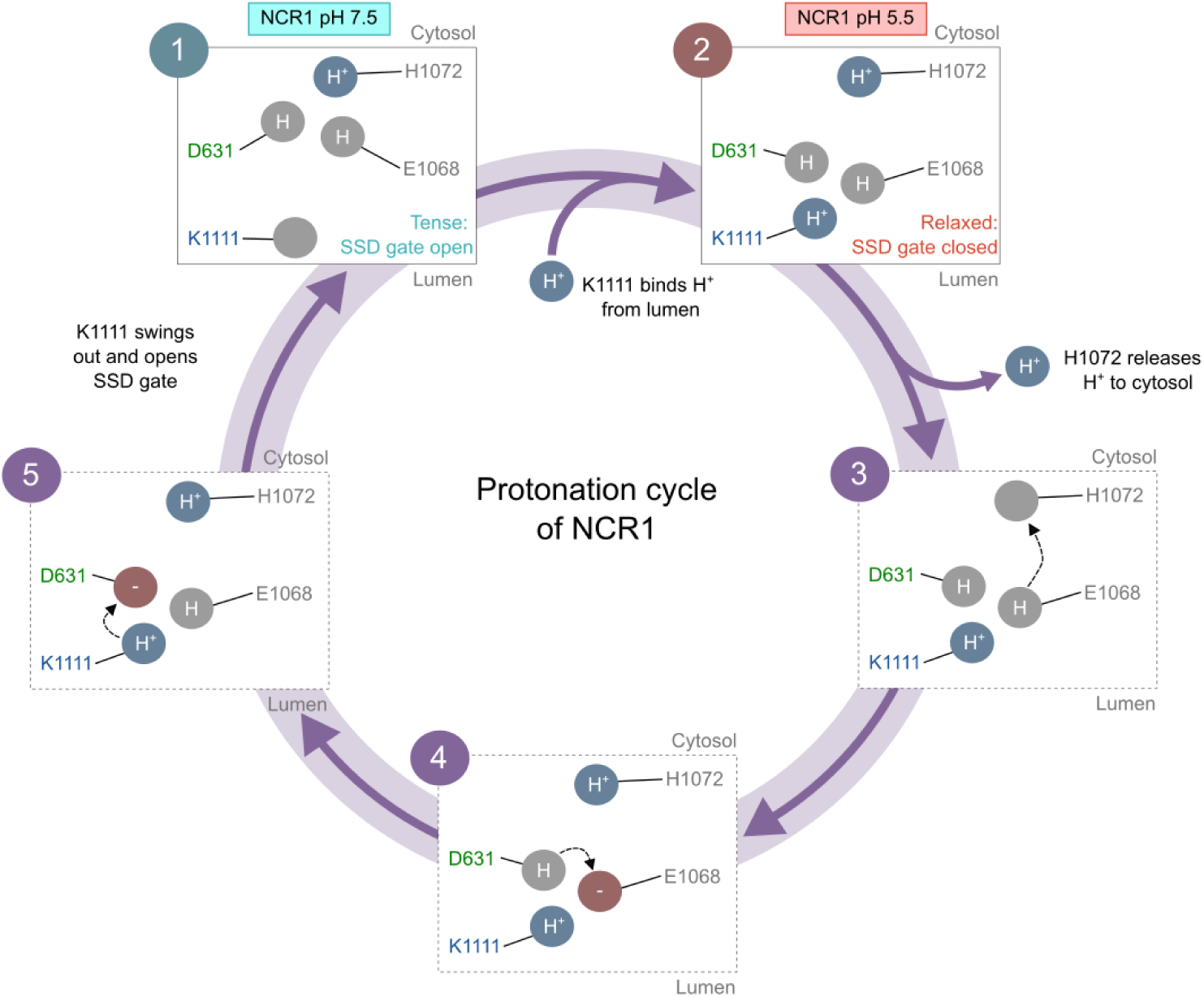
Cartoon representation of the protonation cycle proposed for NCR1. The protonation state of four titratable residues within the transmembrane domains are shown as protonated (blue), neutral (grey) and deprotonated (red). The re-arrangement of charges is shown in five steps: **(1)** Asp631 and Glu1068 interact with His1072. This residue placement represents the tense NCR1 pH 7.5 conformation, where the SSD gate is open. **(2)** Lys1111 binds a proton from the lumen, moves towards and interacts with Asp631 and Glu1068. This residue placement represents the relaxed NCR1 pH 5.5 conformation, where the SSD gate is closed. **(3)** His1072 releases its proton to the cytosol and is subsequently protonated by E1068. **(4)** D631 re-protonates E1068. **(5)** Lys1111 re-protonates Asp631 and then swings away from Asp631 and E1068. The protonation state of the residues is re-established for the next protonation cycle to occur.

The proposed model is consistent with a generalized model for function in the RND superfamily. The transmembrane domain - specifically the symmetry point between the SSD and pSSD - functions as a motor driven by proton binding (or monovalent cation binding for some RND proteins), which drives cycling of the SSD and pSSD between relaxed and tense conformations. This cycle leads to movements of the effector or luminal domains, which in NPC proteins cause the MLD and the CTD to concertedly either contract or relax Zone I and Zone III of the tunnel and propel sterol towards the membrane. A similar switch between a tense and relaxed conformation of the transmembrane domain in the protein Dispatched, another RND protein, is suggested to be central for its function in Sonic Hedgehog release and mobilization (Wang et al., 2021). Also noteworthy, a similar “breathing” motion as we describe here has also been described for the transmembrane domains of the RND protein Patched, and suggested to provide the energy to drive cholesterol egress (Ansell et al., 2023). In conclusion, we have presented evidence for a glycocalyx in the *S. cerevisiae* vacuole and have provided a detailed structural analysis of NCR1 in two distinct conformational states. We have linked changes in tunnel dimensions and in sterol molecule locations to changes of the acidic pair in the transmembrane region. Our model suggests that the conformational transition is dependent on the protonation state of titratable residues in the transmembrane domains, implying that sterol transport is linked to the proton motive force found across the vacuolar/lysosomal membrane. This model is consistent with a general scheme for transport in the RND superfamily and provides novel insights into how NPC proteins have adapted the RND mechanism to move sterols past the glycocalyx and into the lysosomal/vacuolar membrane.

## ACKNOWLEDGMENTS

We acknowledge the EMBION Cryo-EM Facility at iNANO, Aarhus University (application 0397), where data was collected. We also thank eBIC for Cryo-EM data collection (application BI21404). We acknowledge access to the computational infrastructure at the Center for Structural Biology. We express our gratitude to the Bioimaging Core Facility at Aarhus University.

AB was supported by an AUFF-Ukraine Research Fellowship from The Aarhus University Research Foundation and a “Scholars At Risk from Ukrainian Universities” SARU fellowship. DLS was supported by the National Institutes of Health (Grant Agreement No. R35 GM144109). This work was supported by the Danish Council for Independent Research (Grant Agreement No. 0135-00032B), the Carlsberg Foundation (Grant Agreement No. CF19-0127) and the European Research Council (Grant Agreement No. 637372) to BPP.

## AUTHOR CONTRIBUTIONS

Protein expression and purification: KMF, ED, LN and EO.

Cryo-EM sample preparation, data acquisition and image processing: KMF, ED, LN, EO, DLS and BPP.

Model building and analysis: KMF, ED, LN, EO, DLS and BPP.

Vacuole purification and fluorescence microscopy: AB, KMF.

Project conceptualization, design and supervision: DLS and BPP.

Manuscript preparation: KMF, ED, LN, DLS and BPP.

## DECLARATION OF INTERESTS

The authors declare no competing interests.

## ACCESSION NUMBERS

Atomic models have been deposited in the Protein Data Bank (PDB) and Cryo-EM maps have been deposited in the Electron Microscopy Data Bank (EMDB).

NCR1 in GDN at pH 7.5: PDB 8QEB and EMDB EMD-18350.

NCR1 in GDN at pH 5.5: PDB 8QEC and EMDB EMD-18351.

NCR1 in LMNG at pH 5.5: PDB 8QED and EMDB EMD-18352.

NCR1 in peptidisc at pH 7.5: PDB 8QEE and EMDB EMD-18353.

## MATERIALS AND METHODS

### Glycocalyx staining and visualization

Vacuoles were isolated following the protocol outlined previously (Rieder & Emr, 2000). Briefly, BY4741 cells (http://www.euroscarf.de/plasmid_details.php?accno=Y00000) were cultivated in standard Yeast Extract–Peptone–Dextrose (YPD) liquid media at 30 °C, 150 rpm to OD600 between 0.8-1, harvested by centrifugation at 4000 ×*g* for 5 min at room temperature and, resuspended in 100 mM PIPES/KOH pH 9.4, 10 mM DDM). After incubation at 30°C for 10 min, cells were centrifuged as before and the pellet was gently resuspended in 50 mM potassium phosphate pH 7.5, 0.6 M sorbitol, and 0.16x YPD, supplemented with 2 mg/mL of zymolyase. The resulting spheroplasts were incubated at 30 °C for 25 min and then centrifuged at 4 °C for 5 min at 2500 ×*g*. The pellet was resuspended in 15% Ficoll, 10 mM PIPES/KOH pH 6.8), 200 mM sorbitol, then incubated on ice for 2 minutes followed by 75 s at 30 °C. The resulting cell lysate was transferred to ultracentrifuge tubes and layered with a discontinuous gradient consisting of 8%, 4% and 0% Ficoll (Sigma). After centrifugation at 110,000 ×*g* for 90 min at 4 °C, vacuoles were collected from the interface between the 4% and 0% Ficoll. These vacuoles were then stained with FM4-64 (Thermo Scientific) at a final concentration 2.5 μg/mL for 1 h at 4 °C followed by Concanavalin A conjugated with Alexa Fluor 350 (Thermo Scientific) at final concentration 0.5 mg/mL for 3 h at 4 °C. Visualization of both stains was conducted with a Zeiss Axio Observer Z1 microscope equipped with a 63x objective under oil immersion.

### Homologous expression of NCR1 and membrane preparation

Protein expression of codon-optimised NCR1 from *S. cerevisiae* (UniProt: Q12200) was performed as described previously (Winkler et al., 2019). For expression, *S. cerevisiae* strain DSY-5 was transformed with an expression vector based on p423_GAL1 (Mumberg et al., 1994) carrying full length *ncr1* with a C-terminal deca-histidine tag and grown to high density in a bioreactor and harvested after a 22 h induction using galactose (Lyons et al., 2016). Cells were harvested by centrifugation at 2,500 x*g* for 10 min at 4 °C and the pellets were stored at −70 °C.

Cells were homogenized with a beadbeater (Biospec Products) by mixing ∼200 g of cells with 750 mL lysis-buffer (600 mM NaCl, 100 mM Tris-HCl pH 7.5, 1.2 mM PMSF) in pre-chilled metal canisters containing 0.5 mm glass beads (Biospec Products) and using 5 cycles of agitation (1 min on and 2 min off). Cell lysate was filtered, re-supplied with 1.2 mM PMSF and cell debris was removed by centrifugation at 17,000 x*g* for 20 min at 4 °C. The supernatant was centrifuged at 200,000 x*g* for 2 h at 4 °C and the resulting membrane pellets were divided into 3 g aliquots and stored in 500 mM NaCl, 50 mM Tris-HCl, pH 7.5, 20 % (v/v) glycerol at −20 °C.

### Purification of NCR1

Two aliquots of membranes (∼6 g) were solubilized in basis buffer (500 mM NaCl, 50 mM Tris-HCl, pH 7.5, 10 % (v/v) glycerol, supplemented with 1 % (w/v) cholesteryl hemisuccinate (CHS from Anatrace), 0.6 % (v/v) *n*-dodecyl-β-D-maltoside (DDM from Inalco Pharmaceuticals) and 1.6 mg/mL iodoacetamide (Sigma-Aldrich) for 20 min at 4 °C. The sample was sonicated, filtered with 5 μm as well as 1.2 μm pore size syringe filters (Sartorius), and supplemented with 20 mM imidazole prior to loading onto a 5 mL HisTrap HP column (Cytiva) equilibrated with 20 mM imidazole and 0.017 % DDM in basis buffer. The column was washed with 50 mL of basis buffer with 70 mM imidazole and 0.017 % DDM followed by 45 ml of G20 buffer (200 mM NaCl, 20 mM imidazole) supplemented either with 0.01-0.05 % glyco-diosgenin (GDN from Antrance) or 0.025% LMNG, with pH of either 5.5 (20 mM MES) or 7.5 (20 mM Tris-HCl). Finally, 175 units of thrombin were added (Avantor) to cleave the His-tag and 5 mL of this buffer was slowly circulated through the column at 4 °C overnight. The protein was eluted with 15 mL of the corresponding buffer containing 40 mM imidazole. After concentration, the sample was loaded onto a Superdex200 10/300 GL Increase column (GE Healthcare), equilibrated with 200 mM NaCl, containing either 0.01-0.05 % GDN at pH 5.5 (20 mM MES) or pH 7.5 (20 mM Tris-HCl) or 0.005 % LMNG at pH 5.5 (20 mM MES). Peak fractions were pooled, concentrated to ∼9 mg/mL, and immediately used to prepare EM grids. Purification in peptidisc involved additional wash steps on the 5 mL HisTrap HP column (Carlson et al., 2018). After the 50 mL wash with 70 mM imidazole, the sample was further washed with 50 mL of 100 mM NaCl, 50 mM Tris-HCl pH 7.5, 0,008 % of DDM without imidazole followed by 45 ml of peptidisc buffer (100 mM NaCl, 50 mM Tris-HCl, pH 7.5, 0.5 mg/mL peptidisc from GenScript). 5 mL of this buffer were circulated for 30 min at 4 °C followed by a final wash with 100 mM NaCl, 50 mM Tris-HCl pH 7.5. This solution was supplemented with thrombin and circulated overnight, as described earlier. Size-exclusion chromatography was performed using 100 mM NaCl, 10 mM Tris-HCl, pH 7.5 and the peak fraction of the monomer was concentrated to ∼4 mg/mL.

### Cryo-EM sample preparation, imaging and image processing

Samples were applied to glow discharged C-flat 1.2/1.3 μm Cu grids (Protochips, #M-CF313-100) and plunge-frozen in liquid ethane using a Vitrobot Mark IV (Thermo Scientific) at 4 °C and 100 % humidity.

Images for NCR1-75 and NCR1-PD were collected at the Danish National cryo-EM facility, EMBION, at Aarhus University in Denmark at 130,000x magnification on a K3 detector (Gatan) with a pixel size of 0.647 Å and total dose of 60 e^-^/Å^2^. Images for NCR1-55 and NCR1-LMNG-55 were collected at the National Electron Bio-imaging Centre (eBIC), Oxford, United Kingdom 165,000x magnification on a K3 detector with a calibrated pixel size of 0.536 Å and a total dose of 60 e ^-^/Å^2^. Defocus generally ranged from −0.6 to −2.5 μm for all four datasets.

Image processing workflows for each dataset are illustrated in Figs. S2-S5. All processing steps were conducted in cryo-SPARC v3.3.1 (Punjani et al., 2017). The workflows involved standard jobs of patch-based motion correction, CTF estimation, curation to exclude poor quality images with CTF-estimated resolution > 4 Å, high astigmatism, and thick ice. Templates were created from an initial set of particles and used for automated particle picking followed by two rounds of 2D classification using a 160 Å circular mask. The best classes were combined for multiple rounds of *ab-initio* reconstruction with two classes to eliminate damaged particles, empty micelles, and other artifacts. The resulting structures were used as references for multiple rounds of hetero-refinement, which typically separated particles with and without an NTD present. Once a homogeneous set of particles was obtained, a final non-uniform refinement was performed to produce the final structure. Resolution was assessed using both the gold-standard FSC curve and a local resolution job.

### Model building and refinement

A summary of the refinement statistics from all 4 models are shown in Table S1. The crystal structure of NCR1 (PDB: 6R4L) was used as a template for model building. To start, this model was rigid-body fitted to each cryo-EM density map using Chimera (Pettersen et al., 2004) and then subjected to molecular dynamics flexible fitting (MDFF) (Trabuco et al., 2008) using Namdinator (Kidmose et al., 2019). For NCR1-PD, the NTD domain and the M1 helix were removed. For NCR1-LMNG-55, the final NCR1-55 model was initially docked into the density. Using these initial models, COOT v0.9.3 (Casanal et al., 2020; Emsley et al., 2010) was used for manual adjustment of the polypeptide, addition of carbohydrates (NAG) at specific Asn residues (123, 401, 513, 900 and 940 depending on the model) and placement of ligands. These ligands included cholesteryl hemisuccinate (Y01), glyco-diosgenin (Q7G), phosphatidylethanolamine (PTY), and ergosterol (ERG) (Moriarty et al., 2009). The models were gradually improved by iterative rounds of real-space refinement using PHENIX (Adams et al., 2010; Afonine et al., 2018) and manual adjustments in COOT. The final models were validated with MolProbity (Chen et al., 2010), CaBLAM (Prisant et al., 2020) and Ramachandran-Z analysis (Rama-Z) (Sobolev et al., 2020) and the PDB validation server (Gore et al., 2017).

### Model analysis and figure preparation

Caver Analyst 2.0 Beta (Jurcik et al., 2018) was used to identify and analyze the tunnels in NCR1 structures. Caver Analyst 2.0 Beta default parameters were used except that the shell radius was set to 9, the probe radius to 0.2; the searches were initiated at a point between Asp557 and Tyr1130. Structural figures were prepared with USCF Chimera v1.15 (Pettersen et al., 2004) and USCF ChimeraX v1.3 (Pettersen et al., 2021). SEC curves and sterol tunnel dimensions were plotted with GraphPad Prism v9 and incorporated into Inkscape. The multiple sequence alignment of RND protein members was made in PROMALS3D (Pei et al., 2008) and visualized in Aline (Bond & Schuttelkopf, 2009). ImageJ (Schindelin et al., 2012) was used to view and adjust the contrast of micrographs from isolated vacuoles.

## Supplementary Figures and Tables

**Figure S1.**
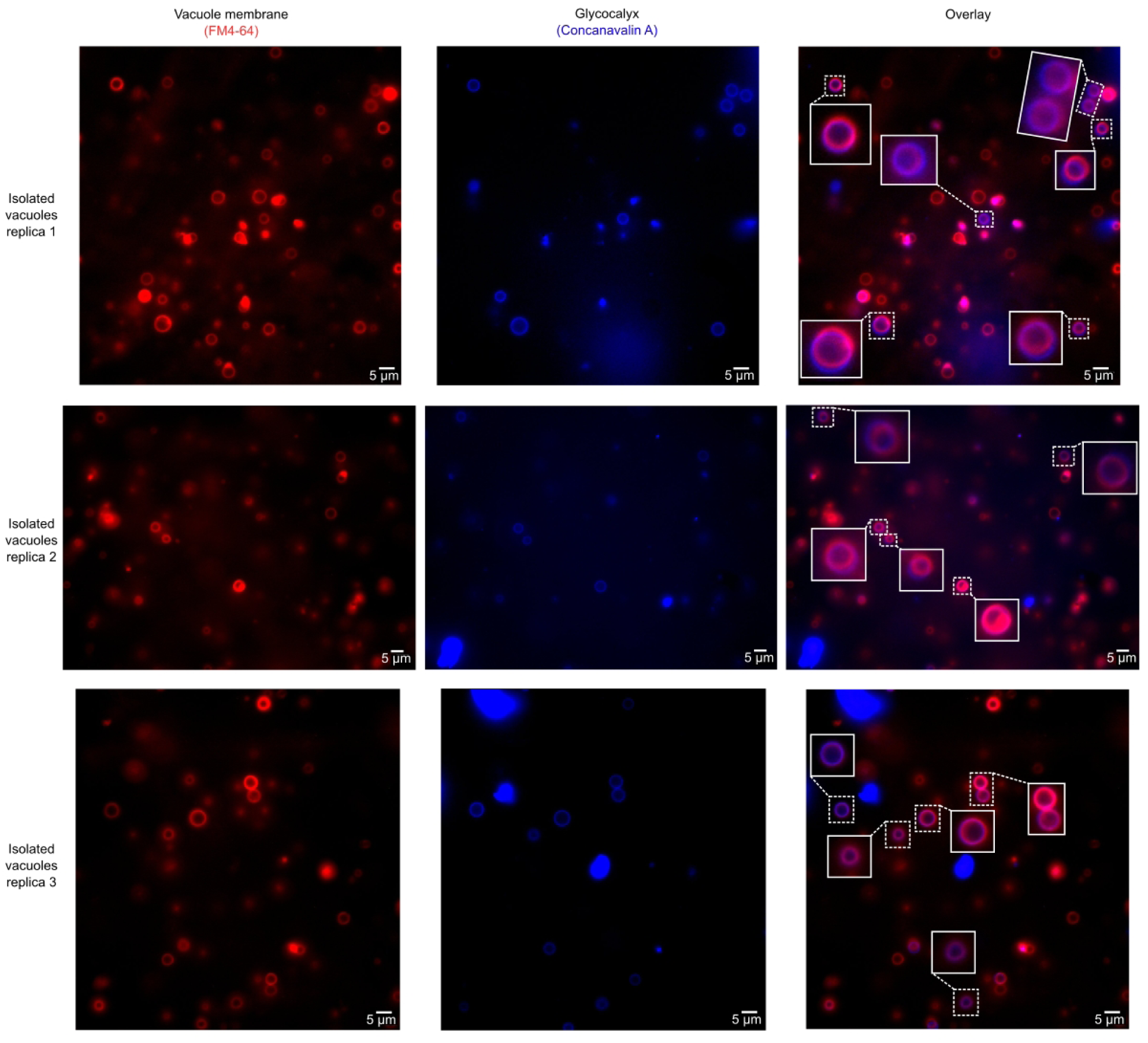
Raw micrographs of co-stained, isolated vacuoles. Vacuolar membranes were stained red with the lipophilic dye FM4-64 and the glycocalyx was stained blue with Concanavalin A conjugated to an AlexaFluor. Co-localization of the dyes shows the glycocalyx appearing immediately inside the vacuolar membrane, producing a purple hue. Isolated vacuoles showing co-localisation of staining are enlarged in boxes with white, solid borders.

**Figure S2.**
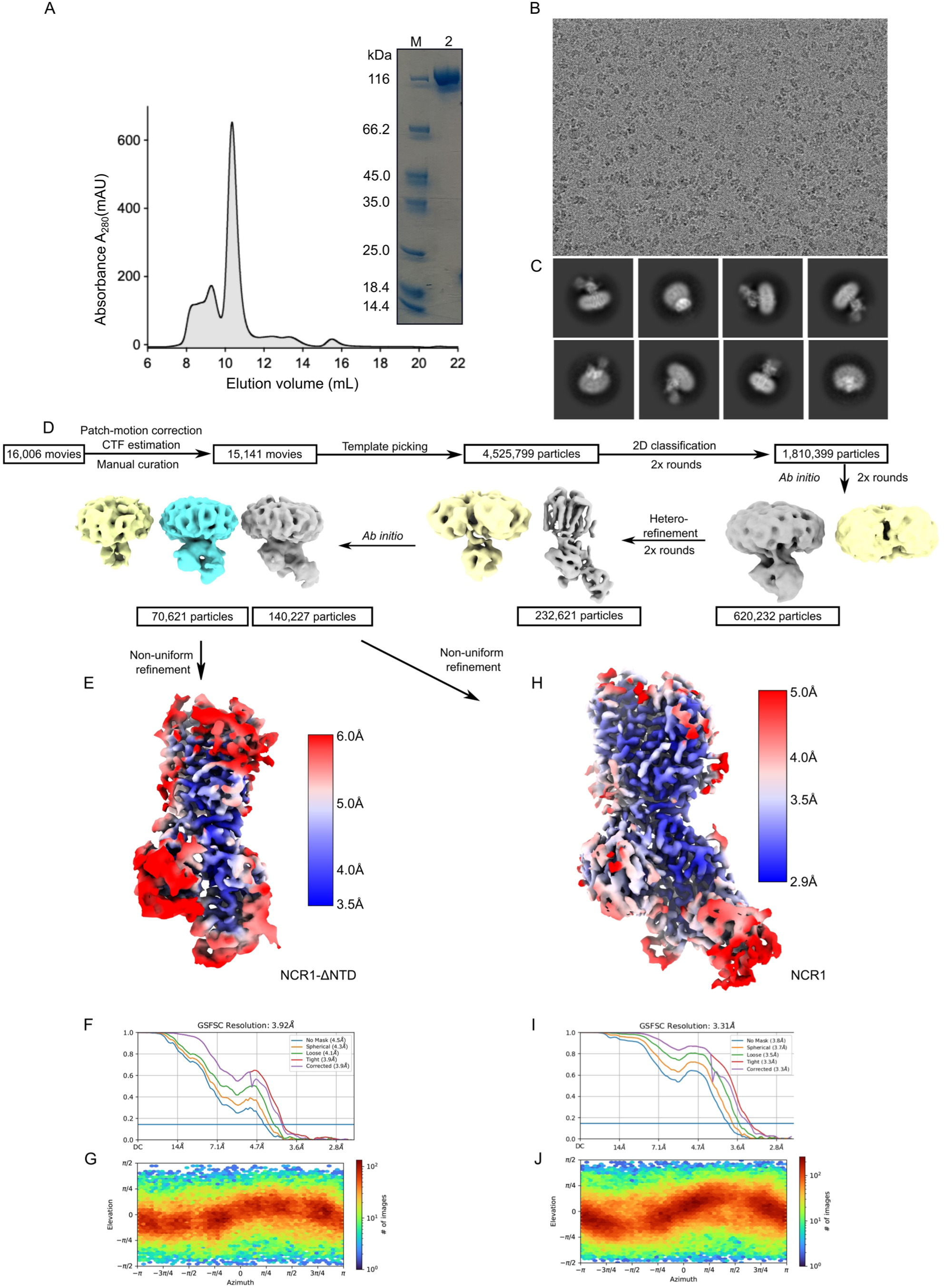
Cryo-EM data processing workflow for NCR1 in GDN at pH 7.5. **(A)** Size-exclusion chromatography (SEC) profile of NCR1 with the peak fraction ∼10.5 mL. The peak fraction was visualized by SDS-PAGE (insert, lane 2). **(B)** Representative motion-and CTF-corrected micrograph. **(C)** Selected 2D classes revealed that NCR1 adopted a variety of orientations. **(D)** Flowchart of the data processing strategy used to separate the empty micelles and degraded particles (yellow) from intact NCR1. Classes for complete NCR1 are shown in grey and a ΔNTD class is shown in cyan. **(E)** Local resolution map for the ΔNTD map colored according to the resolution range shown by the insert. **(F)** Fourier shell correlation (FSC) curve of NCR1-ΔNTD data with resolution at FSC = 0.143 indicated as 3.92 Å. **(G)** Viewing direction distribution plot of NCR1-ΔNTD. **(H)** Local resolution map for complete NCR1 colored according to the resolution range shown by the insert. **(I)** Fourier shell correlation (FSC) curve of complete NCR1 with resolution at FSC = 0.143 indicated as 3.31 Å. **J)** Viewing direction distribution plot of complete NCR1.

**Figure S3.**
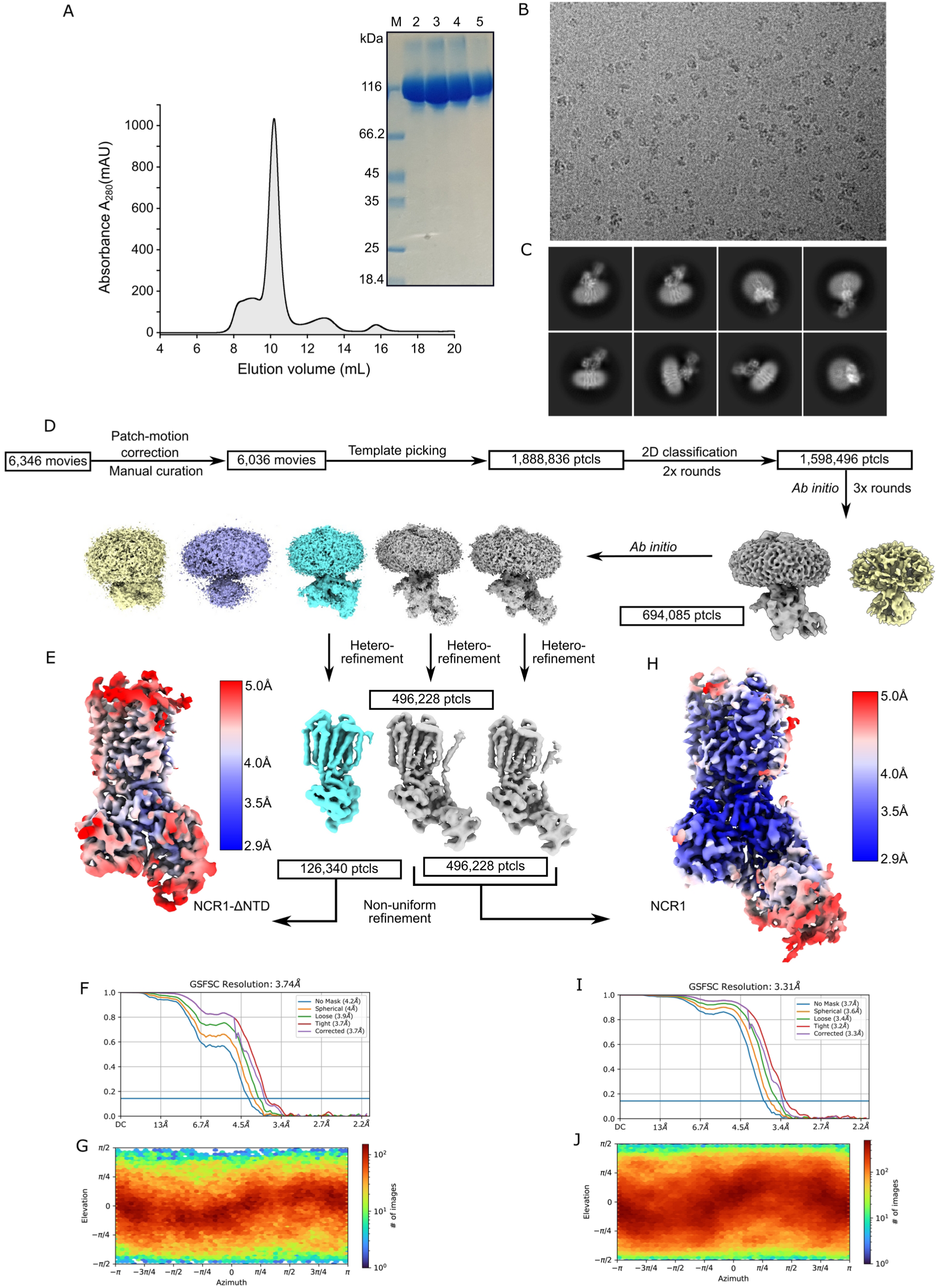
Cryo-EM data processing workflow for NCR1 in GDN at pH 5.5. **(A)** Size-exclusion chromatography (SEC) profile of NCR1 with peak fractions from ∼9-11 mL. These fractions were visualized by SDS-PAGE (insert, lanes 2-5). **(B)** Representative motion-and CTF-corrected micrograph. **(C)** Selected 2D classes revealed that NCR1 adopted a variety of orientations. **(D)** Flowchart of the data processing strategy used to separate the empty micelles and degraded particles (yellow and purple) from intact NCR1. Classes for complete NCR1 are shown in grey and a ΔNTD class is shown in cyan. **(E)** Local resolution map for NCR1-ΔNTD colored according to the resolution range shown by the insert. **(F)** Fourier shell correlation (FSC) curve of NCR1-ΔNTD with resolution at FSC = 0.143 indicated as 3.74 Å. **(G)** Viewing direction distribution plot of NCR1-ΔNTD. **(H)** Local resolution map for complete NCR1 colored according to the resolution range shown by the insert. **(I)** Fourier shell correlation (FSC) curve of complete NCR1 with resolution at FSC = 0.143 indicated as 3.31 Å. **(J)** Viewing direction distribution plot of complete NCR1.

**Figure S4.**
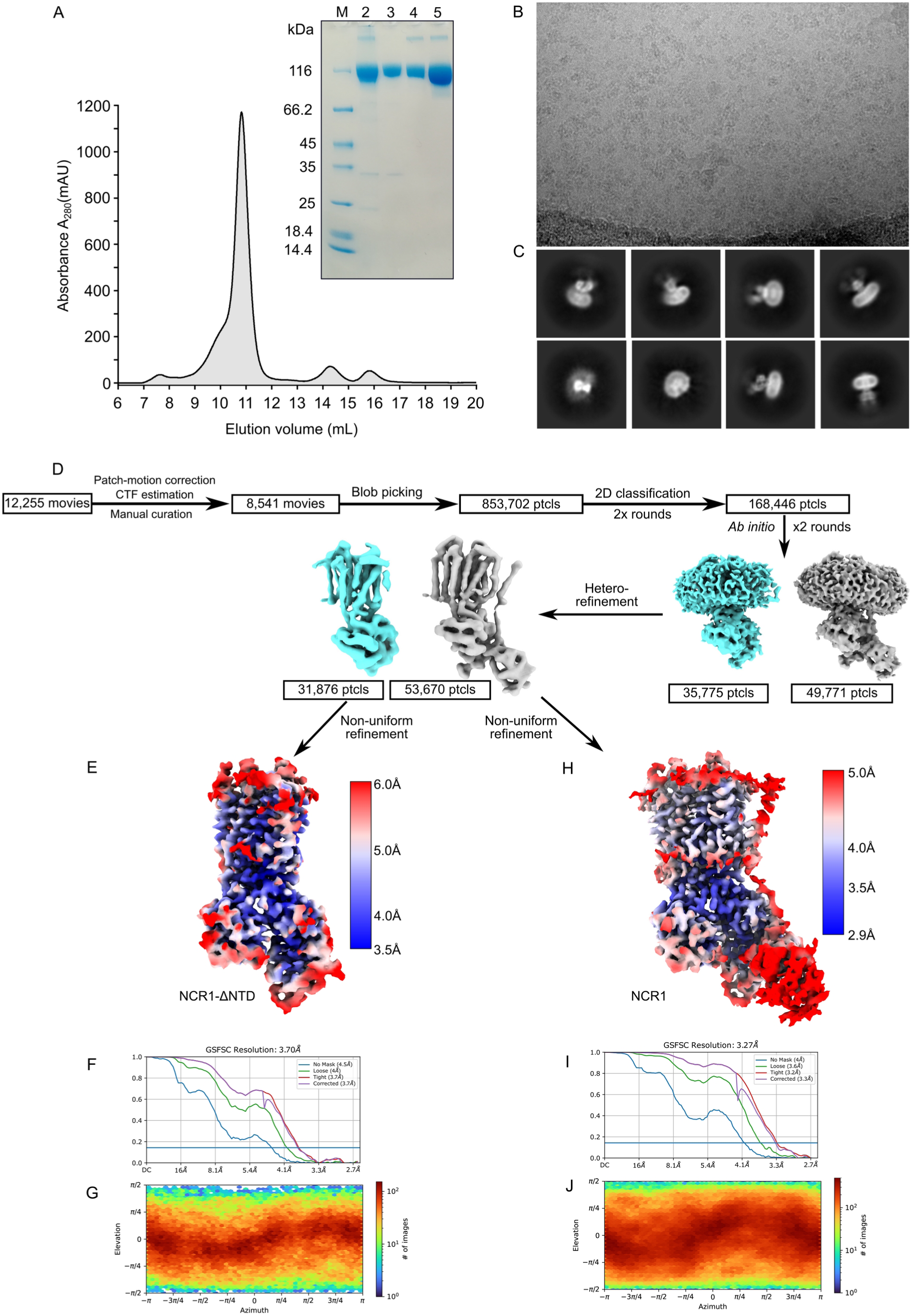
Cryo-EM data processing workflow for NCR1 in LMNG at pH 5.5. **(A)** Size-exclusion chromatography (SEC) profile of NCR1 in LMNG with peak fractions from ∼10.5-11.5 mL. The SDS-PAGE (insert) shows the sample before filtering (lane 2), the sample after filtering (lane 3), SEC peak fraction 1 (lane 4), SEC peak fraction 2 (lane 5). **(B)** Representative motion-and CTF-corrected micrograph. **(C)** Selected 2D classes revealed that NCR1 adopted a variety of orientations. **(D)** Flowchart of the data processing strategy used to separate classes for complete NCR1 (grey) and a ΔNTD class (cyan). **(E)** Local resolution map for NCR1-ΔNTD colored according to the resolution range shown by the insert. **(F)** Fourier shell correlation (FSC) curve of NCR1-ΔNTD with resolution at FSC = 0.143 indicated as 3.70 Å. **(G)** Viewing direction distribution plot of NCR1-ΔNTD. **(H)** Local resolution map for complete NCR1 colored according to the resolution range shown by the insert. **(I)** Fourier shell correlation (FSC) curve of complete NCR1 with resolution at FSC = 0.143 indicated as 3.27 Å. **(J)** Viewing direction distribution plot of complete NCR1.

**Figure S5.**
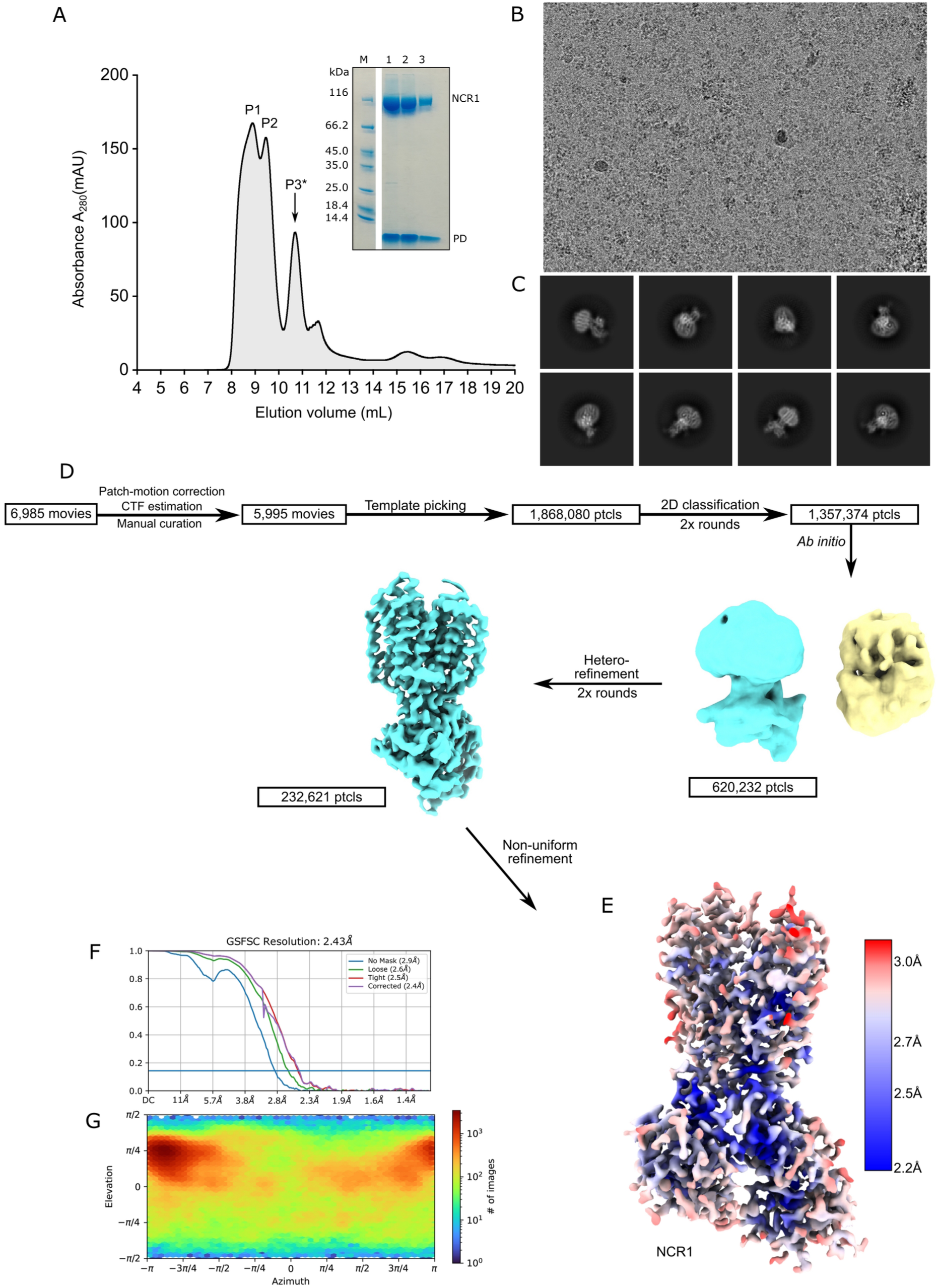
Cryo-EM data processing workflow for NCR1 in peptidisc. **(A)** Size-exclusion chromatography (SEC) profile of NCR1 showing a multimer peak (P1), a dimer peak (P2) and monomer peak (P3*). All three peaks were visualized by SDS-PAGE (insert) where lane 1 corresponds to P1, lane 2 to P2 and lane 3 to P3*. The fraction used for preparing grids is indicated with *. **(B)** Representative motion-and CTF-corrected micrograph. **(C)** Selected 2D classes revealed that NCR1 adopted a variety of orientations. **(D)** Flowchart of the data processing strategy used to separate degraded particles (yellow) from intact NCR1 (cyan). No NCR1 classes showed density for the NTD or the associated M1 helix. **(E)** Local resolution map for NCR1 colored according to the resolution range shown by the insert. **(F)** Fourier shell correlation (FSC) curve of NCR1 with resolution at FSC = 0.143 indicated as 2.43 Å. **(G)** Viewing direction distribution plot of NCR1.

**Figure S6.**
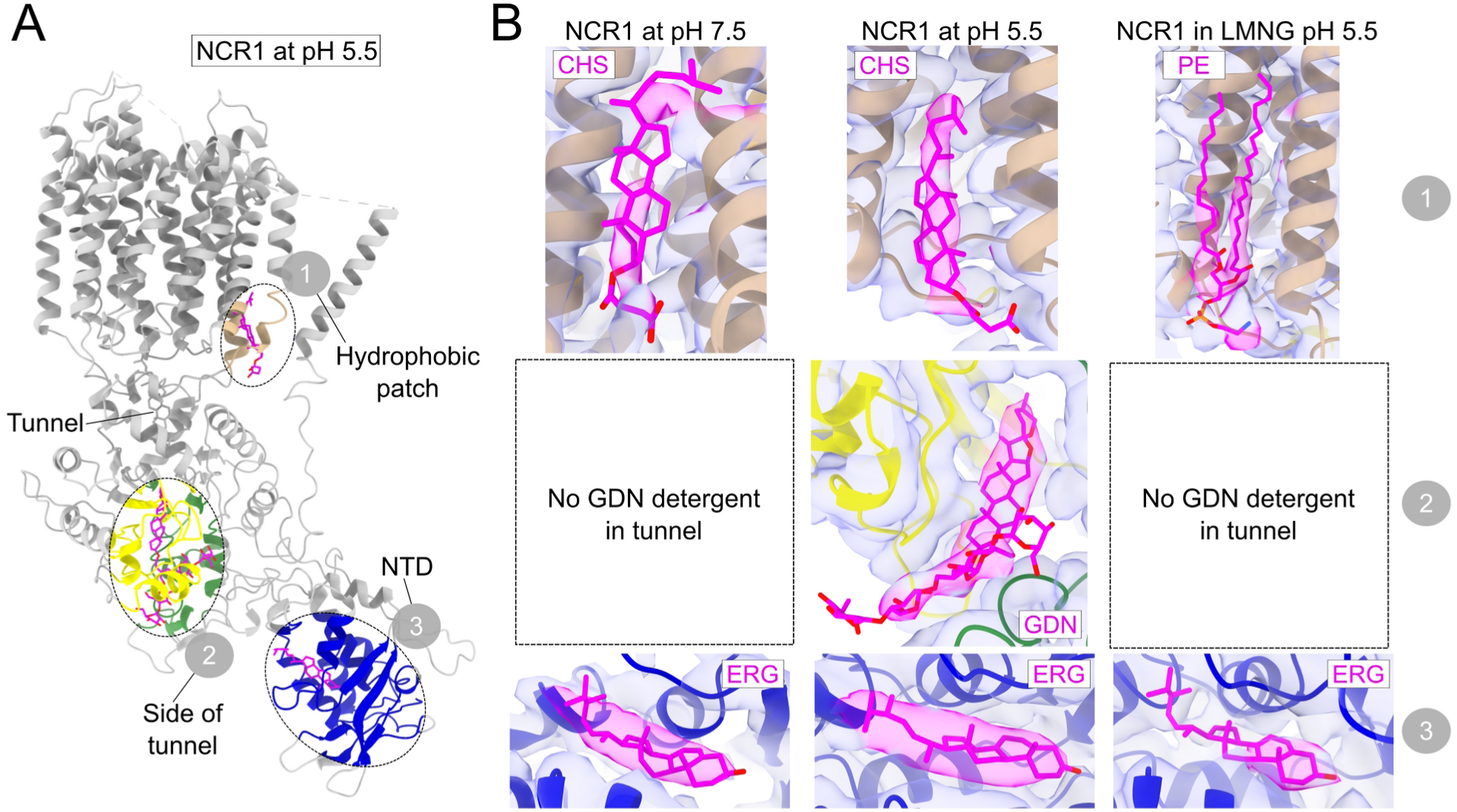
Ligands in addition to sterol inside the tunnel of NCR1 structures solved in detergent. **(A)** Model of NCR1 at pH 5.5 used to highlight regions where ligands are bound. Region 1 is at the hydrophobic patch between the pSSD and M1 helix, region 2 is at the bottom of the tunnel and region 3 is inside the NTD. **(B)** Ligands bound in NCR1 at pH 7.5, NCR1 at pH 5.5 and NCR1 in LMNG at pH 5.5. Only NCR1 at pH 5.5 has a GDN detergent molecule partially within the tunnel.

**Figure S7.**
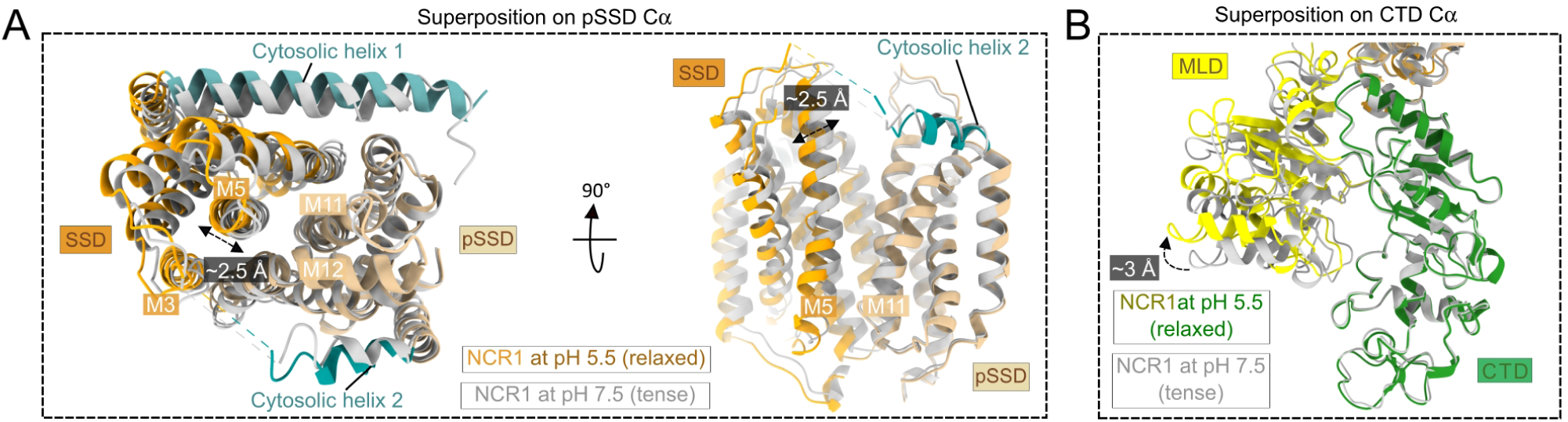
Rigid-body movement of the MLD and CTD. **(A)** Superposition of pSSDs from NCR1 at pH 5.5 (colored) and pH 7.5 (grey) as shown from the perspective of the cytosol (left) and from the front (right). The SSD moves ∼2.5 Å with respect to the pSSD when transitioning between the tense and relaxed conformations, exemplified by helix M5. **(B)** Superposition of CTDs from NCR1 at pH 5.5 (colored) and pH 7.5 (grey) shows a displacement and slight rotation of ∼3 Å of the MLD.

**Figure S8.**
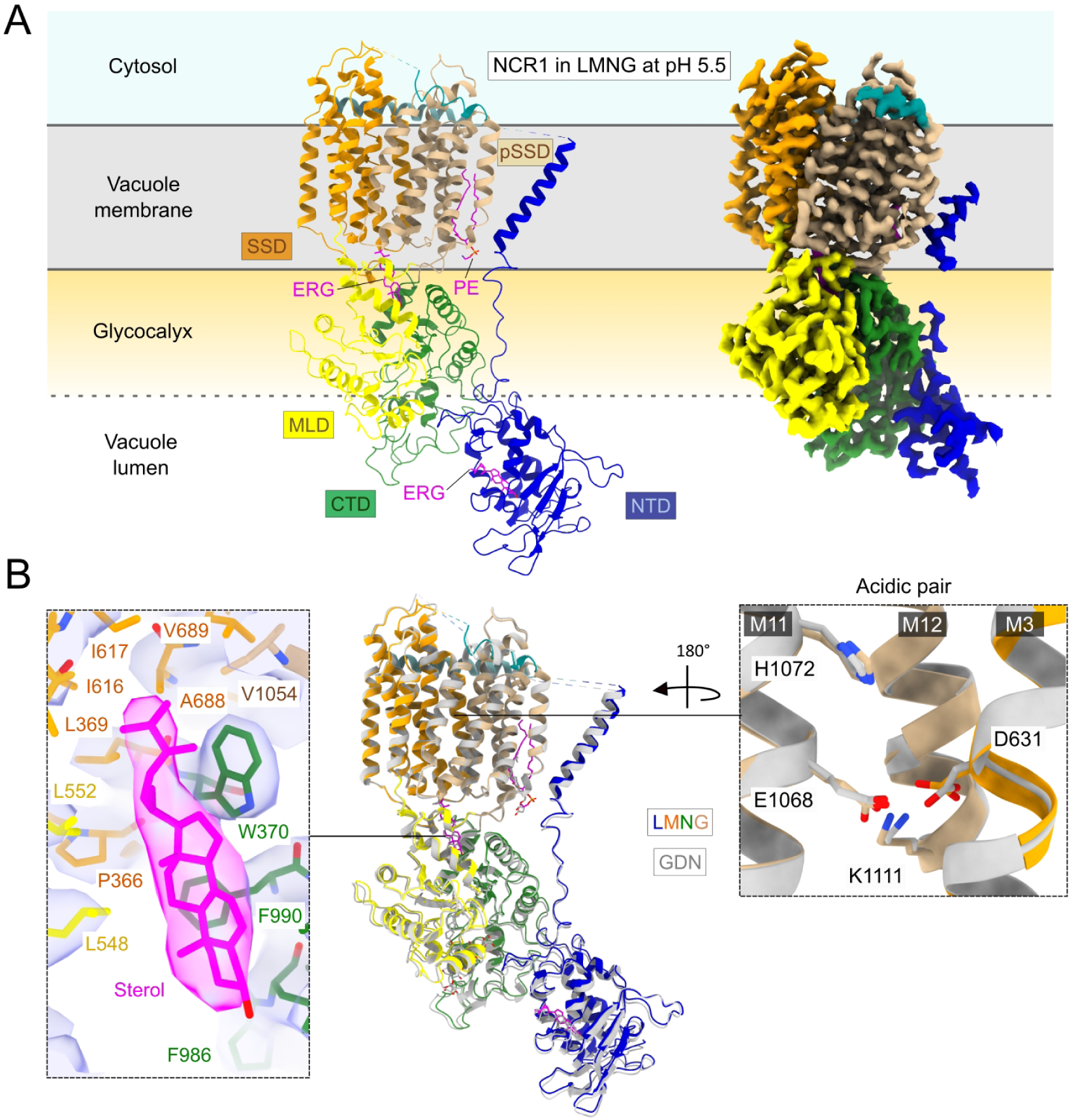
Comparison of NCR1 in GDN and LMNG, both at pH 5.5. **(A)** Model and map of NCR1 in LMNG at pH 55 colored as the topology model. Ligands are depicted in magenta and include an ergosterol in the NTD and the tunnel, and a PE molecule in the hydrophobic patch between the pSSD and M1. **(B)** Density of ergosterol in the tunnel and a structural alignment of NCR1 in LMNG (colors) to NCR1 in GDN (grey), focusing on the acidic pair. The acidic pair interacts with Lys1111 in both structures.

**Figure S9.**
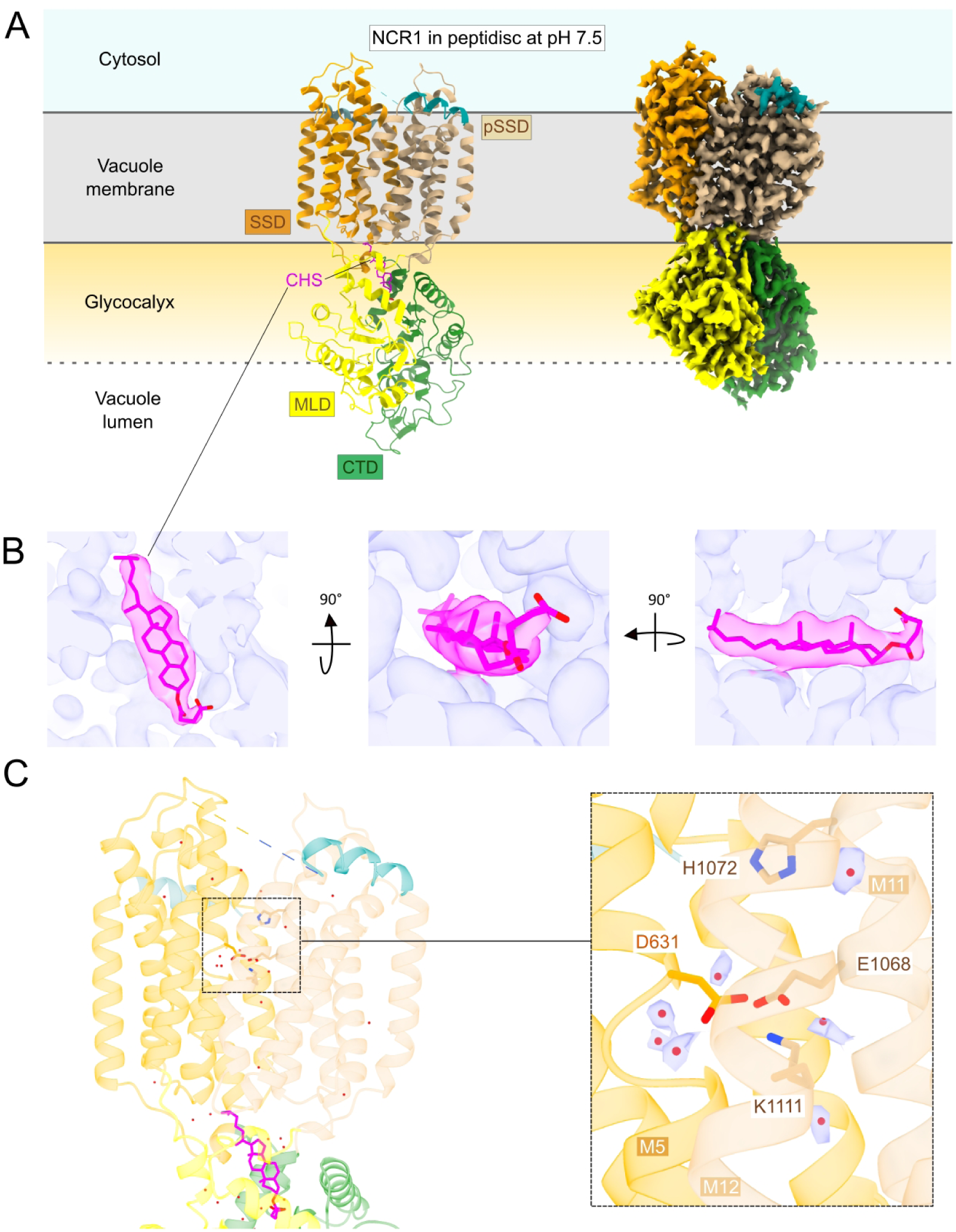
NCR1-PD structure, ligand pose and water densities. **(A)** Map and model of NCR1-PD colored as the topology model. The NTD and M1 are absent and the single CHS ligand in the tunnel is shown in magenta as sticks. **(B)** Different views of the CHS and its surrounding density. **(C)** Densities of water molecules in the vicinity of the acidic pair.

**Figure S10.**
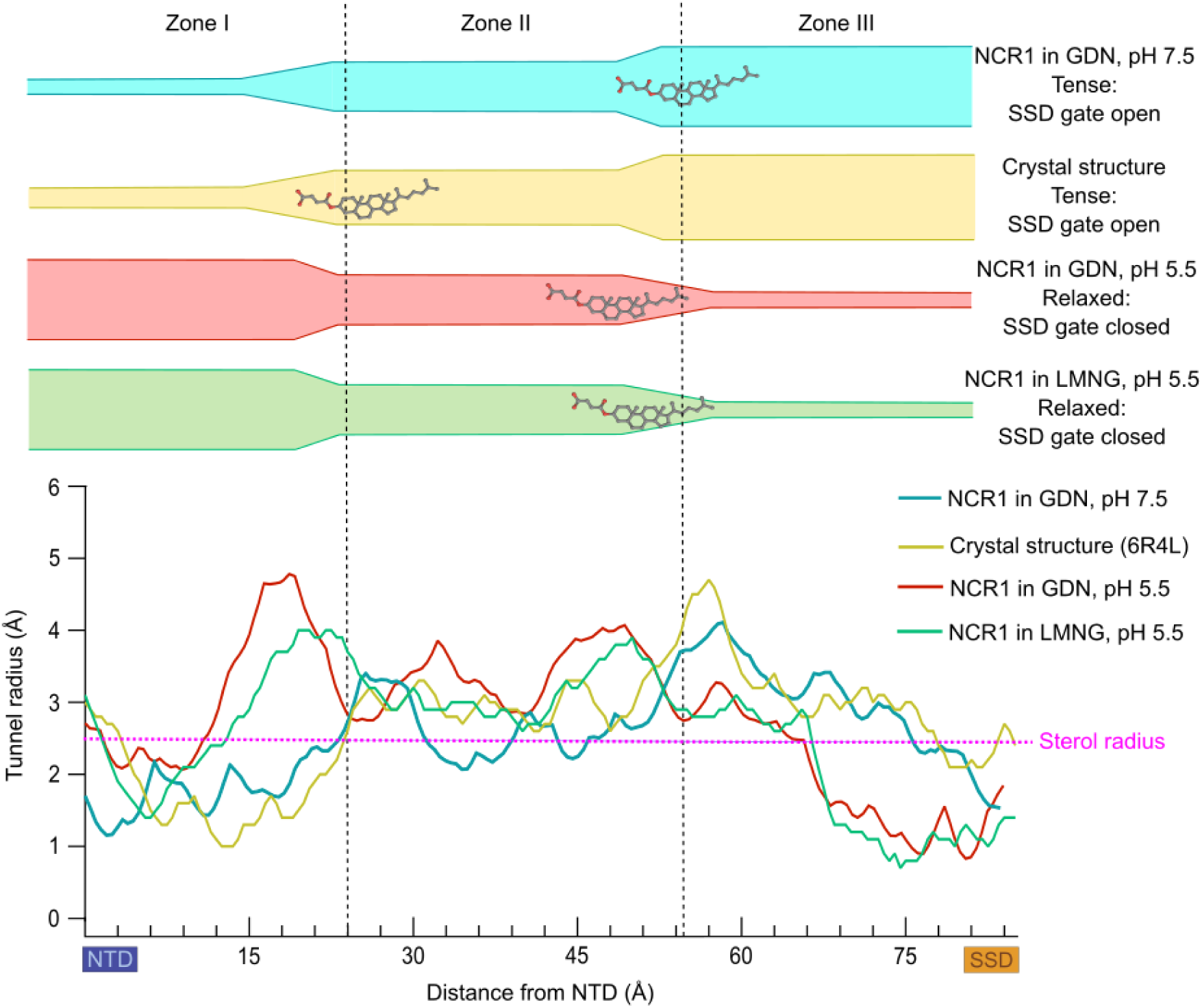
NCR1 tunnel spelunking. Comparison of the tunnel dimensions among the crystal structure and all the structures solved in detergent presented in this paper. The graph indicates the tunnel radius (Å) versus the distance from the NTD (Å). Tunnels depicted include NCR1 at pH 7.5 (cyan), the crystal structure (yellow), NCR1 at pH 5.5 (red) and NCR1 in LMNG at pH 5.5 (green). Sterol radius (2.5 Å) is shown with a dotted magenta line along the X axis. The graph is divided into Zones, I, II and III, which section the schematic representation of the tunnels, along with the sterol location, shown above the graph.

**Figure S11.**
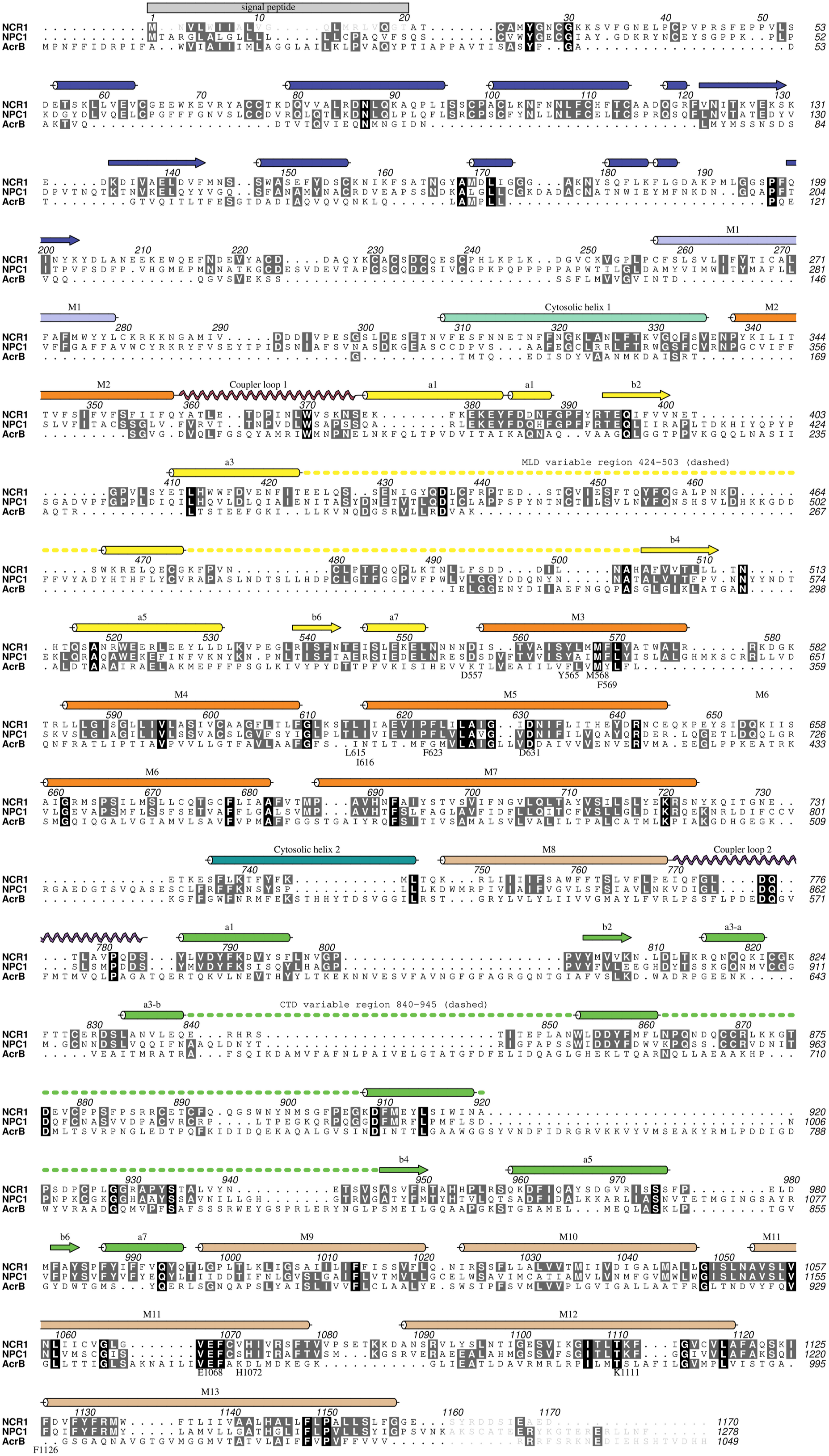
RND protein multiple sequence alignment. Protein sequence alignment of NCR1 (UniProt: Q12200), human NPC1 (UniProt: O15118) and AcrB (UniProt: P31224) performed in PROMALS3D. Secondary structure elements are shown for NCR1.

**Table S1.** Cryo-EM Data Acquisition, Model Refinement and Validation Statistics for NCR1 samples. Summary of data collection, processing and refinement parameters

| Table S1. Cryo-EM Data Acquisition, Model Refinement and Validation Statistics for NCR1 samples |  |  |  |  |
| --- | --- | --- | --- | --- |
| Name | NCR1 at pH 7.5 | NCR1 at pH 5.5 | NCR1 in LMNG at pH 5.5 | NCR1 in peptidisc |
| Detergent | Detergent GDN | Detergent GDN | Detergent LMNG | Peptidisc |
| pH | pH 7.5 | pH 5.5 | pH 5.5 | pH 7.5 |
| Data Collection |  |  |  |  |
| Microscope |  | Titan Krios |  |  |
| EM Facility | EMBION | eBIC | eBIC | EMBION |
| Voltage (kV) |  | 300 |  |  |
| Detector |  | Gatan K3 |  |  |
| Magnification | 130,000 | 165,000 | 130,000 | 130,000 |
| Pixel size (Å/pixel) | 0.647 | 0.536 | 0.651 | 0.85 |
| Electron exposure (e<sup>-</sup>/Å<sup>2</sup>) | 59.558 | 60.000 | 60.000 | 60.600 |
| Defocus range (µm) | 0.2-2.5 | 0.6-2.6 | 0.6-1.8 | 0.4-1.8 |
| Number of movies<sup>#</sup> | 15,141 | 6,036 | 8,541 | 4,223 |
| Reconstruction |  |  |  |  |
| Software |  | cryoSPARC (v3.3.1) |  |  |
| Initial particle images (no.) | 4,525,799 | 1,888,836 | 853,705 | 1,559,630 |
| Final particle images (no.) | 140,227 | 496,228 | 53,670 | 550,587 |
| Symmetry imposed |  | C1 |  |  |
| Map resolution (Å)* | 3.3 | 3.3 | 3.27 | 2.43 |
| Map sharpening B factor (Å<sup>2</sup>) | -108.2 | -179.2 | -73.5 | -72.4 |
| Model Building |  |  |  |  |
| Software |  | Namdinator<br>Coot (v0.9.3) |  |  |
| Refinement |  |  |  |  |
| Software |  | PHENIX (v11.9.2-4158) |  |  |
| Initial model (PDB id) | 6R4L | 6R4L | 6R4L | 6R4L |
| Model composition |  |  |  |  |
| non-hydrogen atoms | 8972 | 9,104 | 9,032 | 6,858 |
| Protein residues | 1098 | 1101 | 1101 | 823 |
| Residue range | 21-1156 | 21-1156 | 21-1156 | 324-1156 |
| Ligands | ERG (ergosterol) = 1<br>YO1 (cholesterol hemmisuccinate) = 2<br>NAG (GlaNAc) = 6 | ERG (ergosterol) = 1<br>YO1 (cholesterol hemisuccinate) = 2<br>Q7G (GDN) = 1<br>NAG (GlaNAc) = 8 | ERG (ergosterol) = 2<br>PTY (PE lipid) = 1<br>NAG (GlaNAc) = 8 | YO1 (cholesterol hemisuccinate) = 1<br>NAG (GlaNAc) = 5 |
| Waters | 1 | 0 | 0 | 130 |
| RMSD |  |  |  |  |
| Bond lenghts (Å) | 0.003 (0 > 4σ) | 0.002 (0 > 4σ) | 0.003 (0 > 4σ) | 0.002 (0 > 4σ) |
| Bond angles (deg) | 0.460 (0 > 4σ) | 0.552 (10 > 4σ) | 0.466 (16 > 4σ) | 0.420 (0 > 4σ) |
| Validation |  |  |  |  |
| MolProbity score | 1.58 | 1.49 | 1.55 | 1.44 |
| Clashscore | 5.31 | 4.26 | 4.27 | 4.01 |
| Rotamer outliers (%) | 0.41 | 0.10 | 0.10 | 1.61 |
| Rama-Z score (Whole/Helix/Loop) | -0.18 / 0.35 / -0.46 | 0.19 / 0.74 / -0.41 | 0.44 / 1.19 / -0.56 | 2.41 / 1.73 / 1.53 |
| CaBLAM outliers | 2.49 | 1.84 | 2.11 | 0.49 |
| Ramachandran Plot Statistics |  |  |  |  |
| Favored regions | 95.70 | 95.89 | 95.07 | 98.29 |
| Allowed regions | 4.30 | 4.02 | 4.75 | 1.71 |
| Disallowed regions | 0.00 | 0.09 | 0.18 | 0.00 |
| Deposited model (PDB id) | 8QEB | 8QEC | 8QED | 8QEE |
| Deposited map (EMDB id) | EMD-18350 | EMD-18351 | EMD-18352 | EMD-18353 |

\*Gold standard FSC with threshold of 0.14. <sup>#</sup>Remaining after manual curation of exposures.

## Notes

### Competing Interest Statement

The authors have declared no competing interest.

